# Discovery of a regulatory node that coordinates cell envelope assembly in mycobacteria

**DOI:** 10.64898/2026.07.31.741936

**Authors:** Ruby Hao Sun, Yushu Chen, Shu-Sin Chng

## Abstract

Mycobacteria possess a complex double-membrane cell envelope critical for survival and pathogenesis. Proper assembly of this architecture requires the biosynthesis and transport of major components, including arabinogalactan (AG) polysaccharides and mycolic acids (MAs), but how these processes are effectively coordinated is unknown. Here, we discover an essential membrane complex that serves as a regulatory node in mycobacterial envelope biogenesis. The acyltransferase TmaT and the arabinofuranosyltransferase AftD physically interact; cryo-EM structures reveal a 1:1 stoichiometry, and present a novel fold for TmaT, featuring a central channel that binds co-factor for acetylation in the periplasm. We establish that the TmaT-AftD interaction, and the catalytic activities of both enzymes, are required for MA transport across the cell envelope, as well as AG ligation to the cell wall, the final stage of AG biosynthesis. The TmaT-AftD complex coordinates the two major envelope assembly pathways, presenting a structural vulnerability for future anti-mycobacterial drug development.

## Introduction

*Mycobacterium tuberculosis*, the causative agent of tuberculosis (TB), and other non-tuberculous mycobacterial (NTM) species, such as *Mycobacterium abscessus*, are well-known human pathogens that are difficult to eradicate^1,2^, in part due to the presence of a complex, multi-layered cell envelope structure that offers a formidable barrier against host defenses and conventional antibiotics. Specifically, mycobacterial cells assemble an outer membrane (OM), rich in long-chain mycolic acids (MAs) that are physically linked to the peptidoglycan cell wall via arabinogalactan (AG) polysaccharides^3,4^. This integrated architecture, collectively known as the mycolyl-arabinogalactan-peptidoglycan (mAGP) complex, is indispensable for mycobacterial survival and pathogenicity, and underpins resistance to external stresses^5,6^.

To assemble this essential mAGP framework, cells initiate AG biosynthesis at the inner membrane (IM) (Figure 1). A linear galactan domain is first polymerized on a lipid carrier by galactosyltransferases^7^, forming the base for the construction of three highly branched arabinan domains by arabinofuranosyltransferases (Ara*f*Ts)^8^. Using decaprenylphosphoryl-D-arabinose (DPA) as the sugar donor, AftA adds the first ara*f* onto the galactan core^9^, and EmbA/B elongates the arabinan chain via linear α−(1→5) linkages^10,11^. AftC then mediates α−(1→3) glycosylation to generate branched arabinan^12,13^, which are terminally capped by AftB using β−(1→2) linkages to provide attachment sites for MAs^14–16^. Another enzyme AftD has been shown to possess α−(1→5) and/or α−(1→3) glycosylation activity, but its exact role(s) in AG biosynthesis remains elusive^17,18^. Once fully assembled in the periplasm, this mature AG precursor is covalently ligated to the peptidoglycan by the ligase Lcp1^19^, followed by the final step of MA transfer to terminal ara*f*s by Ag85 enzymes at the OM to complete the mAGP structure^20^.

**Figure 1.**
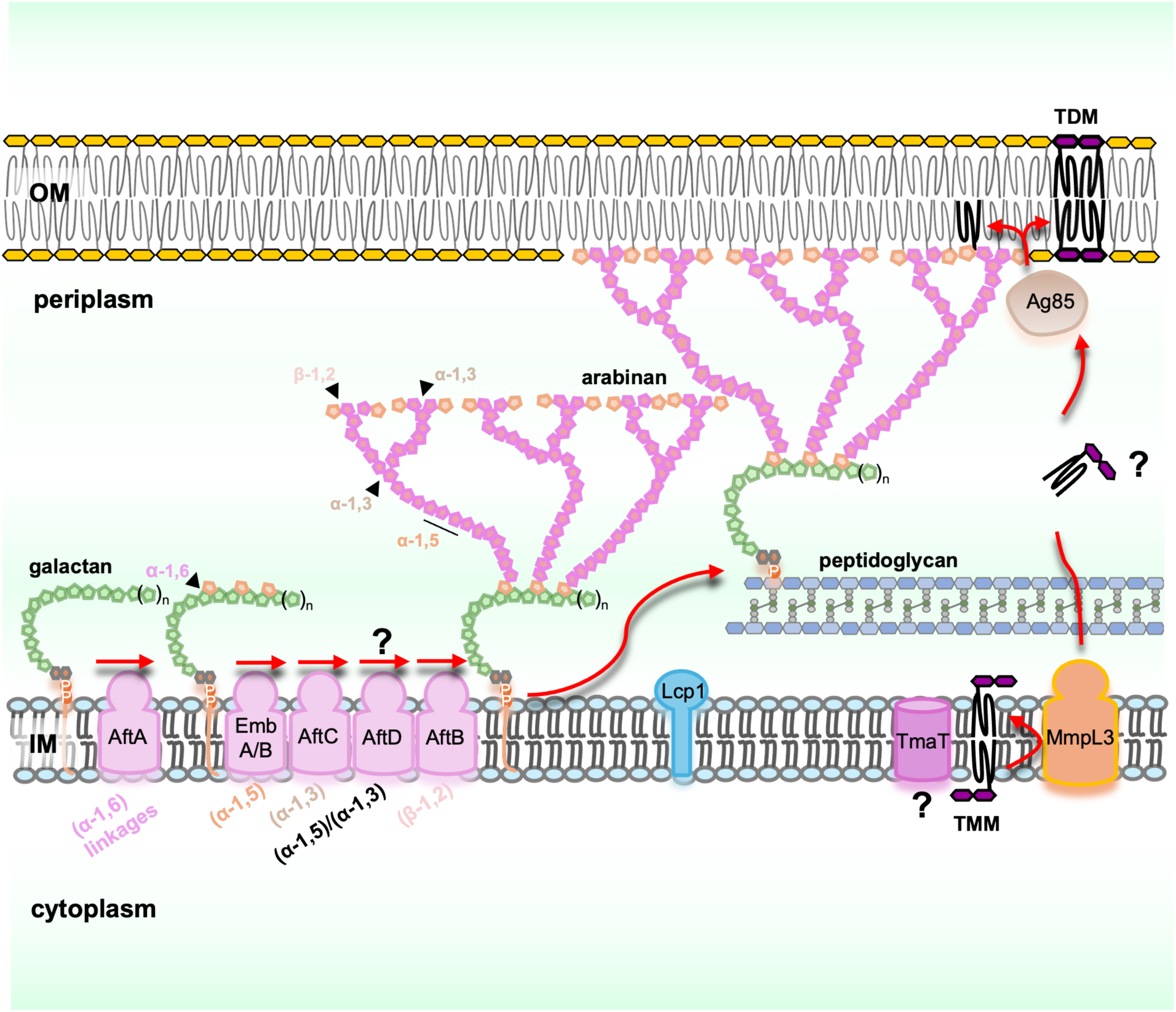
Schematic overview of the assembly of the mycolyl-arabinogalactan-peptidoglycan architecture. mAGP biosynthesis involves the assembly of AG polysaccharides (and ligation onto cell wall), the transport of mycolic acids to the OM, and the mycolylation of the AG termini, as described in the text.

MAs are very long (C_60_-C_90_) α-alkyl, β-hydroxy fatty acids first synthesized in the form of trehalose monomycolates (TMMs) at the IM^21–24^. TMMs need to be transported across the cell envelope in parallel, and coordinated with AG biosynthesis for mAGP assembly (Figure 1). However, much less is known about TMM transport. The essential root-nodulation-division (RND) family transporter, MmpL3, along with its interacting partner TtfA, are believed to flip and/or release TMM across/from the IM^25–28^. Other mycobacterial proteins implicated in TMM transport include Rv0227c/MSMEG_0317^29,30^ and TmaT^31,32^, the latter belonging to the structurally uncharacterized Acyltransferase 3 (AT3) integral membrane protein family; in *Corynebacterium glutamicum*, the TmaT homolog is thought to modify TMM via acetylation to facilitate transport across the IM^31^. Eventually, TMMs are shuttled to the OM via an unknown mechanism as substrates for Ag85 enzymes, which transfer the mycolate moieties onto AG to form mAGP, or onto another TMM to produce trehalose dimycolate (TDM)^20^. How trans-envelope TMM transport is orchestrated is unclear, but it must necessarily be coordinated with AG biosynthesis for optimal mAGP assembly. How the mycobacterial cell synchronizes these two essential envelope pathways also remains to be elucidated.

Here, we report the discovery of an integral membrane complex that is required for both MA transport and AG biosynthesis, the two major branches of mycobacterial cell envelope assembly. We establish that TmaT and AftD form a specific, stable complex, and elucidate cryo-EM structures of this complex in both apo and co-factor-bound states. The structures reveal an extensive TmaT-AftD interface, and a central channel in TmaT that positions acetyl-CoA for acetylation in the periplasm. Our functional analyses demonstrate that both the active sites of TmaT and AftD, as well as their physical association, are required for TMM transport and the ligation of AG to the peptidoglycan scaffold, thus critical for the proper assembly of the mAGP architecture. Together, our findings establish the structural basis for a new regulatory node in the biogenesis of the mycobacterial cell envelope, where the coordinated action of TmaT and AftD ensures the synchronized maturation of the protective OM.

## Results

### TmaT forms a stable complex with AftD in *M. smegmatis*

In our efforts to understand the role(s) of mycobacterial proteins implicated in TMM transport, we sought to identify and characterize native interacting partners via affinity purification. When using His_6_-tagged TmaT as the bait protein, we found a distinct protein species specifically co-purified (Figure 2A). His-tagged TmaT (∼47 kDa) migrated anomalously on SDS-PAGE as a multi-pass membrane protein at ∼40 kDa. Tandem mass spectrometry (MS/MS) identified the putative interacting partner (∼150 kDa) as AftD (MSMEG_0359, Table S1), the essential arabinofuranosyltransferase required for the biogenesis of AG^17,18^. We demonstrated that TmaT could be reciprocally pulled down using His_6_-tagged AftD in *M. smegmatis*, validating this interaction (Figure 2B, Table S2). We further showed that AftD and TmaT co-purified when heterologously overexpressed in *Escherichia coli*, indicating the two proteins interact directly (Figure 2C). We conclude that TmaT and AftD interact specifically to form a membrane protein complex in *M. smegmatis*. Interestingly, this complex is also found in *Corynebacterium glutamicum* in a parallel study^33^.

**Figure 2.**
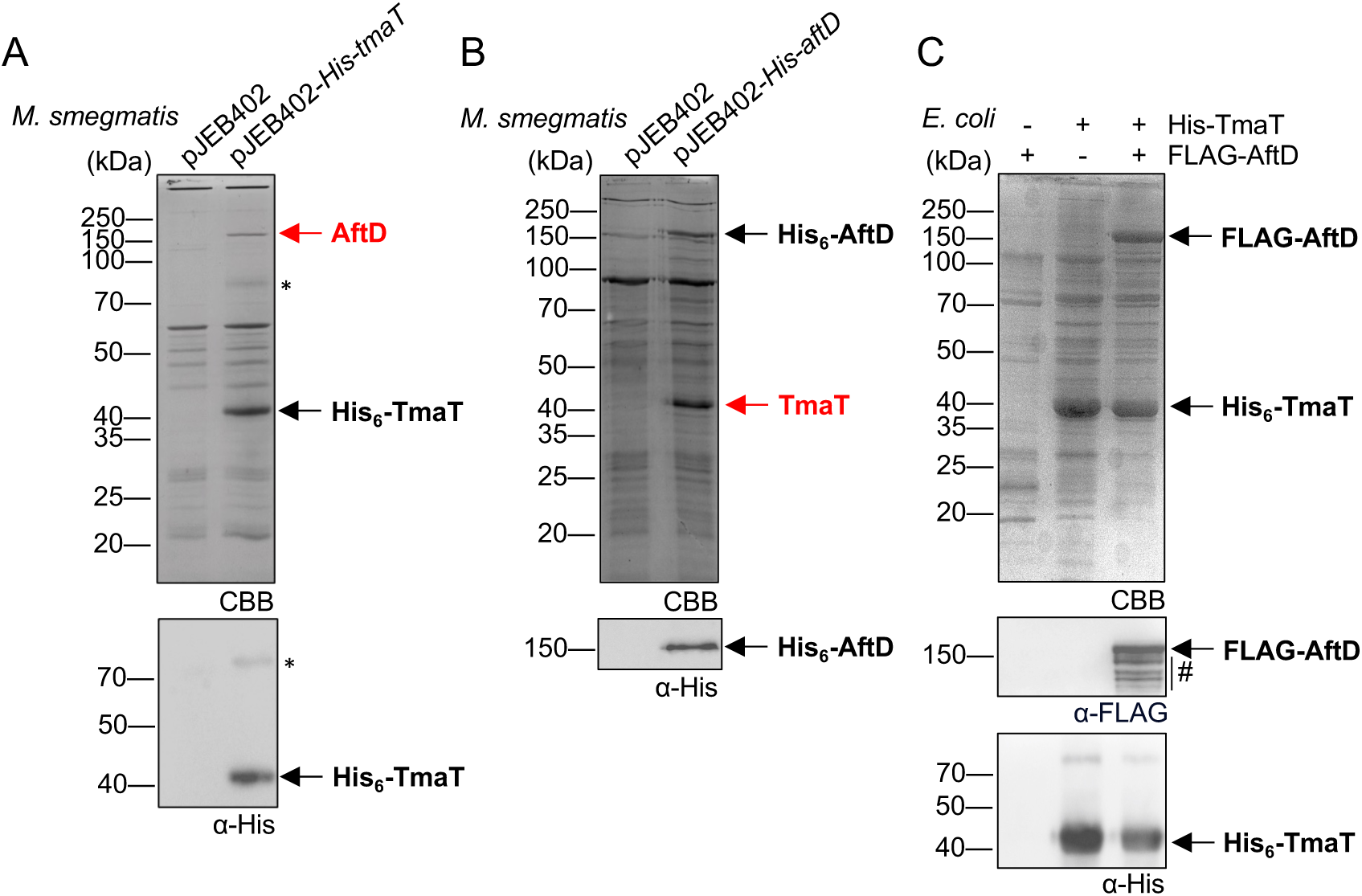
TmaT and AftD interact specifically to form a stable complex. Affinity purification of **(A)** His-tagged TmaT (His_6_-TmaT) or **(B)** His-tagged AftD (His_6_-AftD) from *M. smegmatis* WT cells expressing these proteins from the integrative pJEB402 plasmid. Cells carrying the empty vector serve as negative control. For both experiments, prominent bands specifically co-purified with His-tagged bait were excised and identified by MS. **(C)** Co-affinity purification of recombinant proteins His_6_-TmaT and/or FLAG-tagged AftD overexpressed in *E. coli* cells using pCDFDuet-1 and/or pET22/42 plasmids. For all affinity purification experiments, eluates were analyzed via SDS-PAGE, visualized using Coomassie Brilliant Blue (CBB) staining and α-His/α-FLAG immunoblots, where relevant. Possible aggregation band of TmaT and degradation products of AftD are denoted with * and # respectively.

### Both TmaT and AftD are required for TMM Transport

TmaT and AftD are believed to be separately required for TMM transport and AG biosynthesis^17,18,31,32^, both contributing to the formation of the mAGP complex in the cell envelope. Since the two proteins specifically interact, we hypothesized that they cross-regulate each other. We first asked if AftD also affects TMM transport, like TmaT. To test this idea, we generated *M. smegmatis* conditional *aftD* or *tmaT* knockout (cKO) strains, with an episomal copy of *aftD* (Figure 3A) or *tmaT* (Figure 3B) under an acetamide-inducible promoter, and profiled their extracted lipid compositions by [^14^C]-acetate labeling followed by thin-layer chromatography (TLC). Consistent with previous reports, *tmaT* depletion abolished growth (Figure S1) and TMM transport, as judged by the accumulation of TMM and a corresponding reduction in TDM (Figure 3C and 3D) and MA linked to mAGP (Figure 3E)^32^. Surprisingly, we discovered similar lipid phenotypes in *M. smegmatis* upon *aftD* depletion, which could be rescued by *aftD* expression (Figure 3C-3E), indicating AftD is also required for supporting mycolic acid transport.

**Figure 3.**
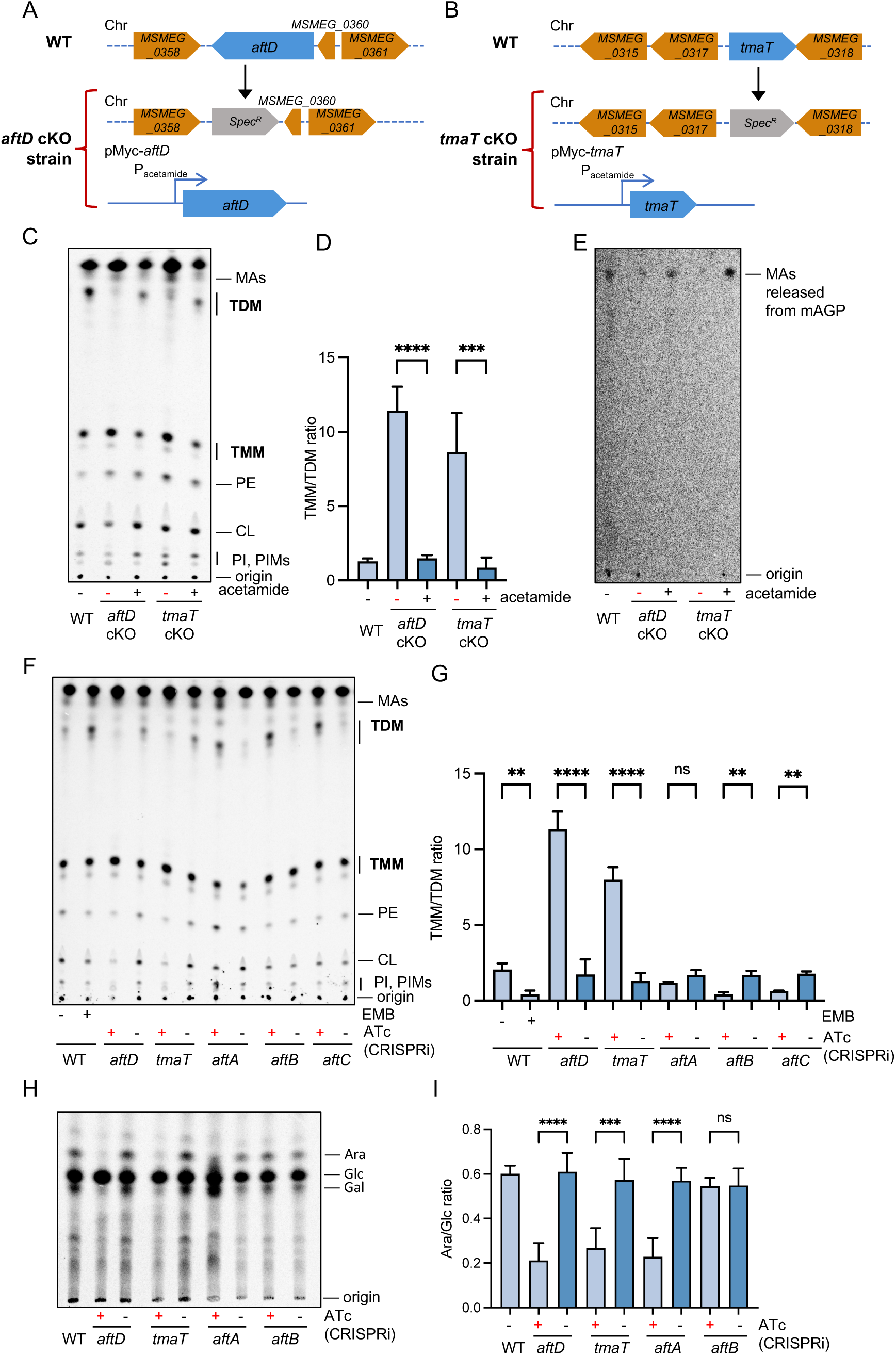
Both TmaT and AftD are required for TMM transport and AG maturation. Genotypes of the **(A)** *aftD* (*MSMEG_0359*), and **(B)** *tmaT* (*MSMEG_0319*) conditional knock out (cKO) strains. See Figure S1 for the corresponding growth curves, where gene expression was induced or not in the presence or absence of acetamide. **(C)** Representative TLC analysis of total extractable lipids isolated from WT, *aftD* cKO, and *tmaT* cKO strains grown in the presence or absence of acetamide. TMM:TDM ratios were quantified and averaged across three independent biological replicates, and shown in **(D)** with error bars representing standard deviations (± SD). **(E)** Representative TLC analysis of MAs released from the mAGP complex in the same strains as (C). **(F)** Representative TLC analysis of total extractable lipids isolated from cells where *aftD*, *tmaT*, *aftA*, *aftB*, or *aftC* expression was knocked down using CRISPRi, or WT cells where EmbABC transferases were inhibited with ethambutol (EMB). For CRISPRi, anhydrotetracycline (ATc) was added to induce inactive Cas9 and the target guide RNA. TMM:TDM ratios were quantified and averaged across three independent biological replicates, and shown in **(G)** with error bars (± SD). **(H)** Representative TLC analyses of [^14^C]-labeled sugars released after hydrolysis from the insoluble cell wall-linked mAGP fractions isolated from WT strains where *aftD*, *tmaT*, *aftA*, or *aftB* expression was knocked down using CRISPRi. Ara:Glc ratios were quantified and averaged across three independent biological replicates, and shown in **(I)** with error bars (± SD). Lipid and sugar TLCs were developed in chloroform/methanol/water (30:8:1, v/v/v) and acetic acid/chloroform/water (7:6:1, v/v/v) solvent systems, respectively. CL, cardiolipin; PE, phosphatidylethanolamine; PI, phosphatidylinositol; PIMs, phosphatidylinositol mannosides; Ara, arabinose; Glc, glucose; Gal, galactose. Student’s *t*-test: ns, not significant; **, *P* < 0.01; ***, *P* < 0.001, ****, *P* < 0.0001. Where a target gene depleted, red (“-” acetamide or “+” ATc) symbols are used.

To assess the specificity of this effect, we investigated other enzymes involved in AG biosynthesis. CRISPRi-mediated knockdown of *aftA*, *aftB*, or *aftC* did not phenocopy *aftD* or *tmaT* depletion, neither did ethambutol-mediated inhibition of the Emb arabinosyltransferases (Figure S2, 3F and 3G)^11,34^. Instead, these perturbations, which block downstream AG biosynthesis, caused a significant accumulation of TDM (Figure 3F and 3G), demonstrating that transport of TMMs to the OM is intact but these substrates are shunted from mAGP to TDM synthesis in the absence of AG attachment sites^20,35^. Taken together, these findings establish a specific role for AftD, distinct from other arabinosyltransferases, in modulating MA transport.

### Both TmaT and AftD are required for AG biosynthesis

We next investigated whether TmaT is reciprocally required for AG biosynthesis, in a manner similar to AftD. Here, we assessed the levels of newly synthesized AG polysaccharides that are attached to the cell wall as part of the mAGP complex, by measuring the amounts of [^14^C]-labeled arabinose and galactose released from insoluble mAGP after hydrolysis. As controls, we observed a drastic reduction of arabinose, but not galactose, in the cell wall upon *aftA* depletion, yet no change to both when *aftB* was depleted, consistent with the functions of AftA and AftB in initiating and terminating the arabinan domain, respectively (Figure 3H and 3I). Intriguingly, we found that the depletion of *aftD* greatly reduced both arabinose and galactose content in the cell wall, suggesting a profound impact on mAGP formation (Figure 3H and 3I). Even more remarkably, *tmaT* depletion closely phenocopied *aftD* depletion, corroborating the idea that TmaT is also required for AG assembly (Figure 3H and 3I).

While AftD has been shown to exhibit α−(1→5) and/or α−(1→3) arabinofuranosyltransferase activity, removing it does not abolish arabinan formation, nor should it impact galactan biosynthesis^17,18^. The loss of AG from the cell wall fraction upon *aftD* depletion indicates that these newly synthesized polysaccharides might be accumulating elsewhere, possibly as lipid-linked precursors in the IM^8^. Indeed, we detected high molecular weight band smears (when analyzed by SDS-PAGE) in the soluble fraction (50% ethanol) accompanied by a marked increase in arabinose levels, upon *aftD* or *tmaT* depletion, fully consistent with the accumulation of lipid-anchored polysaccharides (Figure S3). In comparison, no such changes were observed when *aftA* or *aftB* was depleted (Figure S3). Collectively, these results indicate that in the absence of either TmaT or AftD, lipid-linked AG polysaccharides continue to be synthesized but fail to be efficiently ligated to the peptidoglycan scaffold. Since both proteins are required for TMM transport and AG biosynthesis, we conclude that the TmaT-AftD complex coordinates these two essential envelope processes to ensure proper mAGP assembly.

### Cryo-EM structure reveals the architecture of the TmaT-AftD complex

To gain a molecular basis for its function, we determined the structure of the His_6_-TmaT-AftD complex using single particle cryo-electron microscopy (cryo-EM). The complex was natively purified from *M. smegmatis*, and reconstituted into peptidiscs^36^. After multiple rounds of 2D classification and filtering, 3D reconstruction afforded a density map at ∼4.0 Å nominal resolution (∼20k particles) (Figure S4). Rigid body fitting of an AlphaFold3 model^37^, followed by real space refinement, yielded the structure for the TmaT-AftD complex with a 1:1 stoichiometry (Figure 4A). As expected, our *M. smegmatis* AftD structure is highly similar to the one reported for *M. abscessus* AftD (RMSD 1.24 Å), comprising 16 transmembrane (TM) helices and a large periplasmic domain between TM11’ and TM12’, rich in β-sheets. However, no density corresponding to acyl carrier protein was observed (Figure S5)^38^.

**Figure 4.**
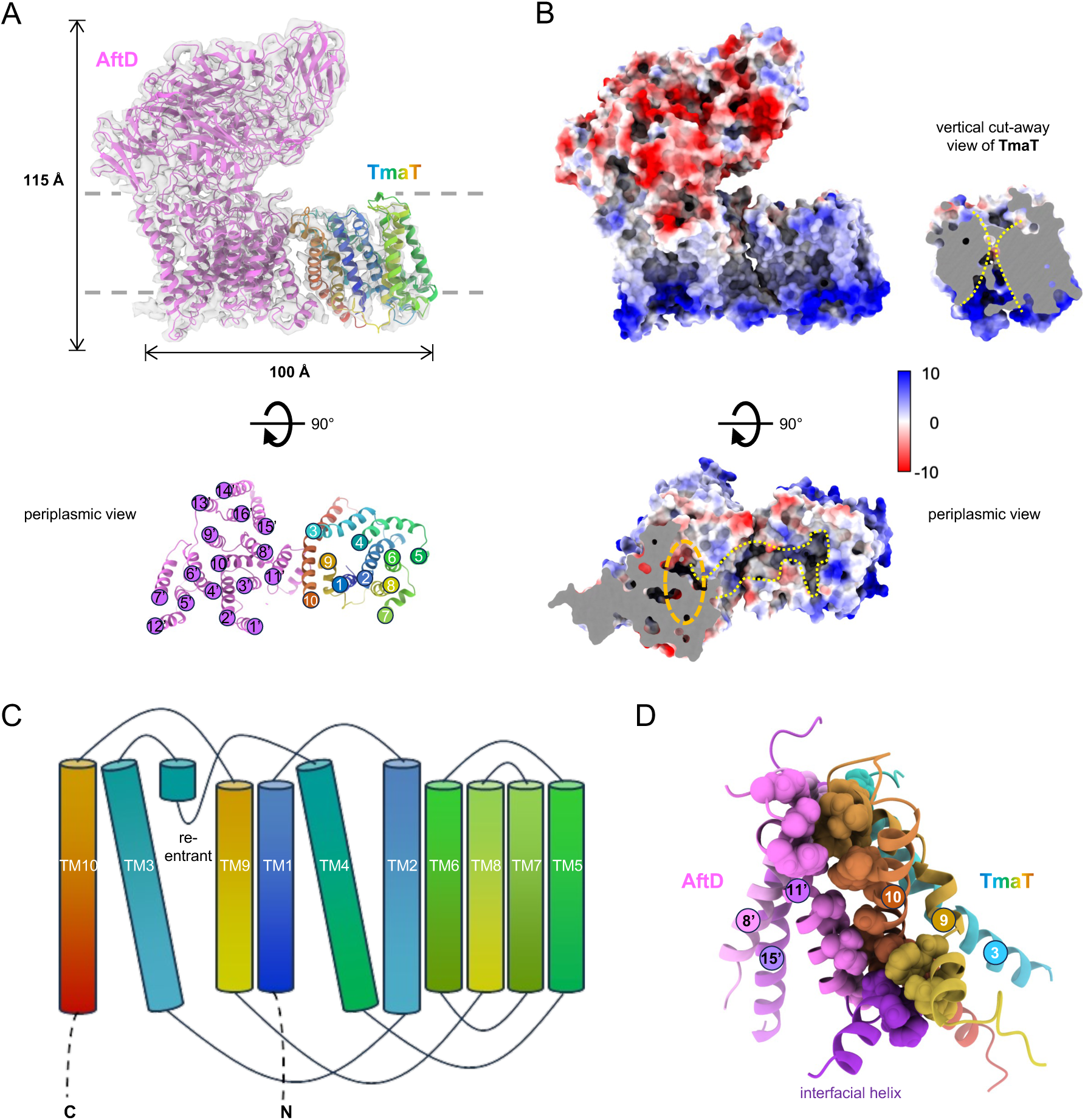
The structure of the 1:1 TmaT-AftD complex reveals an extensive interacting surface, and a novel fold for acyltransferase 3 family protein TmaT. **(A)** Cartoon illustrations of the side view of the TmaT-AftD structure (PDB: **22NM**), showing well-fitted and refined backbones within the EM density map (**EMD-68514**, contour level 0.2, transparency 80%, 4.0 Å resolution). The periplasmic view of the transmembrane region of the complex, revealing TM arrangements of each protein, is shown below. **(B)** Vacuum electrostatic surface representation of the side view of the TmaT-AftD complex, with a cut-away of TmaT revealing the central channel (yellow dashed lines) shown on the right. The periplasmic view of the complex is shown below, with a cut away of periplasmic domain of AftD. A periplasmic groove (yellow dashed outline) connects the TmaT channel to the AftD catalytic site (orange dashed oval). **(C)** 2D topological representation of the TmaT transmembrane domain. **(D)** Cartoon representation of extensive surface area (776 Å^2^, PISA)^44^ between TmaT and AftD. Specific residues in contact are represented by sphere-style atoms.

The structure of TmaT or its AT3 homologs has not previously been experimentally determined; we found that it contains 10 TM helices and a short re-entrant helix, featuring a central channel largely demarcated by TM1, TM2, TM4 and TM9 that widens towards the periplasm (Figure 4B and 4C). TmaT exhibits a distinct fold from other membrane-embedded acyltransferases with known structures, such as the MBOAT proteins^39,40^. The cytoplasmic entrance of the channel is lined by positively charged residues, presumably for binding an acyl-CoA substrate, as predicted for other homologs (Figure 4B)^41^. To determine how the acyl-CoA substrate is bound, we also elucidated the structure of the TmaT-AftD complex incubated with acetyl-CoA, resolved to a nominal resolution of ∼3.4 Å (∼28 k particles) (Figure 5A and S6). Acetyl-CoA could be fitted well into a distinct density observed in the slightly expanded channel (Figure 5A), with the 3’-phosphoryl adenosine diphosphate group occupying the cytoplasmic entrance, interacting with a tetrad of arginines (R27, R96, R99 and R365) through electrostatic interactions and hydrogen bonding (Figure 5B and 5C). Interestingly, the binding pose of acetyl-CoA extends the pantetheine arm towards the periplasmic side, positioning the thioester moiety at the midpoint of the channel, ready for acyl transfer to a periplasmic substrate (Figure 5B and 5C). H38 is part of the conserved R-X_10_-H motif^42,43^, and it sits above the acetyl group near the periplasmic opening of the channel; it may serve as the catalytic base for acetylation of intervening substrate (Figure 5C).

**Figure 5.**
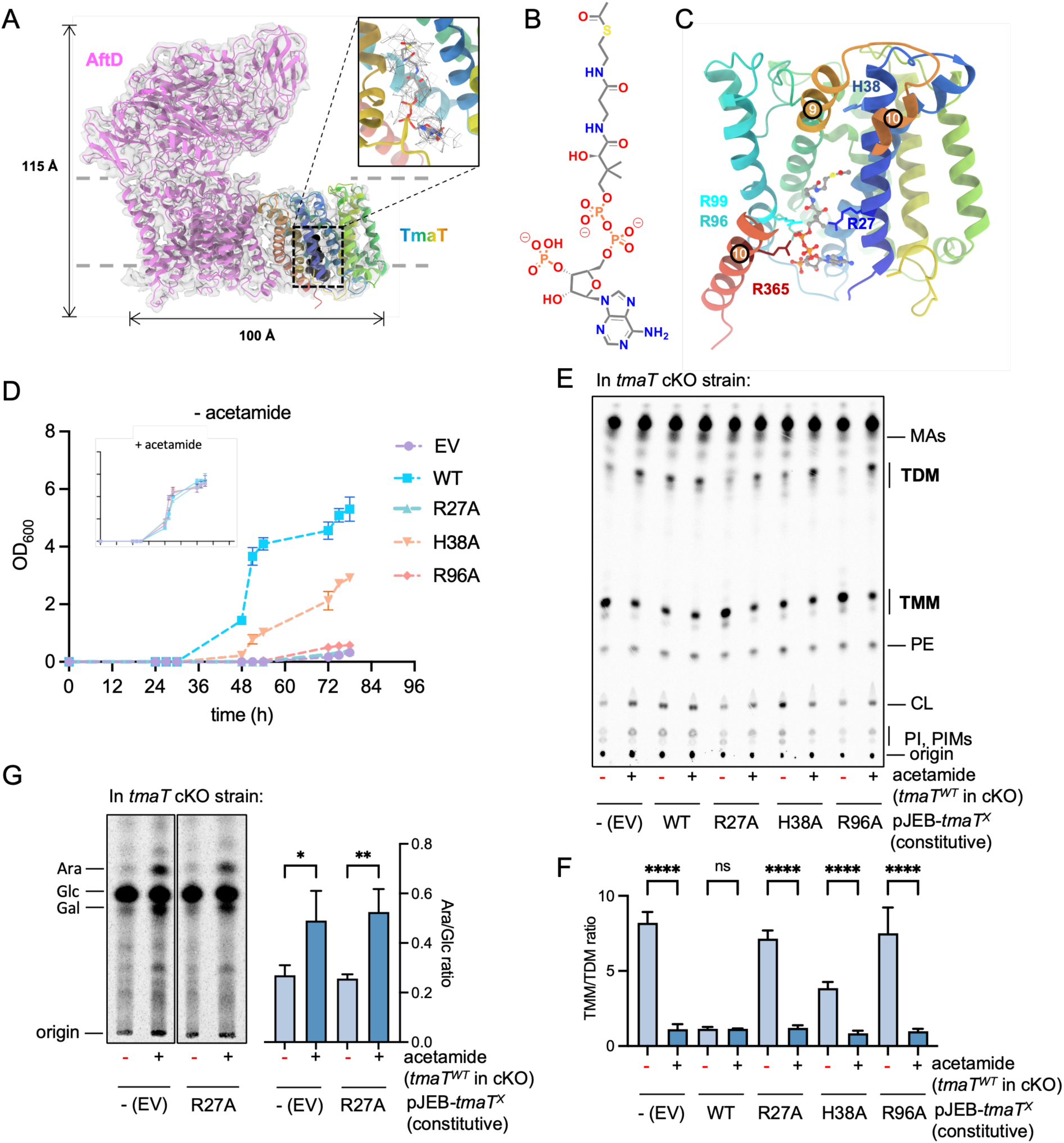
The structure of acetyl-CoA-bound TmaT-AftD complex reveals the catalytic site of TmaT essential for supporting TMM transport and AG biosynthesis. **(A)** Cartoon illustrations of the side view of the structure of TmaT-AftD with acetyl-CoA bound (PDB: **22NS**), showing well-fitted and refined backbones within the EM density map (**EMD-68526**, contour level 0.2, transparency 80%, 3.4 Å resolution). The acetyl-CoA molecule is represented by sphere-style atoms colored in black in the cartoon. A zoom-in view of acetyl-CoA fitted into the density map is shown as an inset (TM1 is removed for better visualization). The density within a 3.5 Å range of the fitted ligand is shown as mesh. **(B)** The chemical structure of acetyl-CoA. **(C)** Cartoon representation of a zoom-in view of TmaT with the bound acetyl-CoA shown as ball and stick. Indicated active residues are shown in sticks. Parts of TM9 and TM10 are hidden to allow better visualization. **(D)** Growth curves of the *tmaT* cKO strain grown in the presence (+) or absence (-) of acetamide to control *tmaT* expression, additionally complemented with an empty vector pJEB402 (EV) or plasmids constitutively expressing WT TmaT or the indicated active site mutants in 7H9 liquid medium. These TmaT variants are stably expressed, and interact well with AftD, comparable to WT TmaT (Figure S7). **(E)** Representative TLC analysis of total extractable lipids isolated from the indicated strains in (D). TMM:TDM ratios were quantified from the TLCs and averaged across three independent biological replicates, and shown in **(F)** with error bars (± SD). **(G)** Representative TLC analyses of [^14^C]-labeled sugars released after hydrolysis from the insoluble cell wall-linked mAGP fractions isolated from *tmaT* cKO strains grown in the presence (+) or absence (-) of acetamide to control *tmaT* expression, additionally complemented with an empty vector pJEB402 (EV) or the plasmid constitutively expressing indicated mutant. Ara:Glc ratios were quantified and averaged across three independent biological replicates, and shown with error bars (± SD). Student‘s t-test: ns, not significant; ****, P < 0.0001. Where a target gene depleted, red (“-” acetamide) symbols are used.

TmaT and AftD interact extensively in the membrane. Across the interface, TM9 and TM10 of TmaT contact TM8’ and TM11’ of AftD, creating a buried hydrophobic surface area of 776 Å^2^ (Figure 4D)^44^. Notably, a short interfacial helix of AftD (before TM11’) contributes to the TmaT-AftD interaction at the cytoplasmic side (Figure 4D). This helix was unresolved in the reported structure for *M. abscessus* AftD alone^38^, but present in our *M. smegmatis* TmaT-AftD structures, suggesting that the interaction with TmaT rigidified this structural motif. Though of unknown significance, the TmaT-AftD interaction essentially connected the TmaT channel to the AftD catalytic domain via a periplasmic groove akin to a substrate binding site (Figure 4B bottom).

### Catalytic activities within the TmaT-AftD complex are critical for its function in coordinating mAGP assembly

We next performed structure-guided mutagenesis to assess whether the catalytic sites within the TmaT-AftD complex are required for its function. We first examined the TmaT active site. Single alanine substitutions at R27/R96 or H38, expected to disrupt CoA binding or impact catalysis in TmaT^41,45^, abolished or impaired growth, respectively (Figure 5D). In addition, we observed defects in TMM transport that correlated with (lack of) growth (Figure 5E and 5F); specifically, the R27A (or R96A) mutation strongly impacted TMM transport. Remarkably, the TmaT^R27A^ variant also caused a severe reduction in AG (arabinose and galactose) linked to the cell wall (Figure 5G), indicating that TmaT-mediated acetylation is a prerequisite not only for MA transport but also for AG incorporation. The active site of TmaT is required for the collective function of the TmaT-AftD complex in coordinating the two pathways.

We then focused on the active site of AftD. Previous structural and functional studies of AftD identified essential and conserved residues in a putative active site^38^. We generated single alanine substitution variants at five positions (K23A, D25A, E235A, K345A, D459A) in *M. smegmatis* AftD, and showed that they failed to complement growth of the *aftD* cKO strain, though stably produced (Figure 6A, S8A and S8C). Remarkably, lipid profiling of cells expressing these active site mutants but depleted of WT *aftD* revealed diminished TMM transport (TMM accumulation and TDM reduction) (Figure 6B and 6C). Furthermore, we tested one of these variants (AftD^K345A^), and found that it cannot support AG incorporation into the cell wall on its own (Figures 6I). All AftD variants could be stably expressed and co-purified with TmaT in *E. coli*, indicating preservation of the TmaT-AftD interaction (Figure S8B). We conclude that the catalytic activity of AftD is necessary for both TMM transport and AG biosynthesis. The catalytic functions of the TmaT-AftD complex ensures proper mAGP assembly.

**Figure 6.**
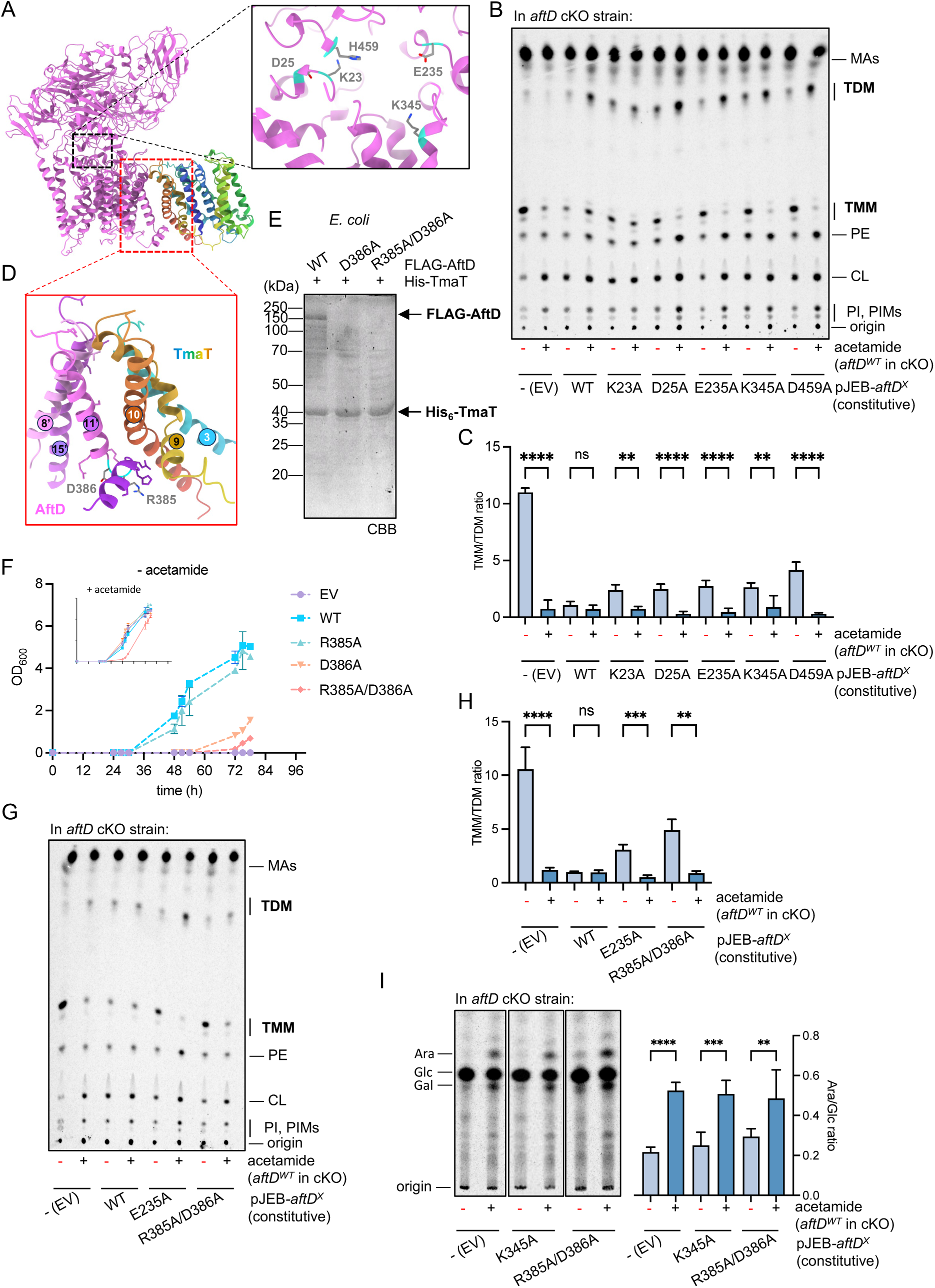
The catalytic activity of AftD and its interaction with TmaT are required for TMM transport and AG assembly. **(A)** Cartoon representation of TmaT-AftD structure with a zoom-in view of the AftD active site shown as an inset. Indicated active residues are shown in sticks with cyan backbones. **(B)** Representative TLC analysis of total extractable lipids isolated from the *aftD* cKO strain grown in the presence (+) or absence (-) of acetamide to control *aftD* expression, additionally complemented with an empty vector pJEB402 (EV) or plasmids constitutively expressing WT AftD or the indicated active site mutants. **(C)** TMM:TDM ratios were quantified from the TLCs in (B) and averaged across three independent biological replicates, and shown with error bars (± SD). **(D)** A zoom-in view of the interface between TmaT and AftD, where indicated residues that disrupted the interaction when substituted are shown in sticks colored by atoms with cyan backbones. Other residues that did not impact TmaT-AftD interaction are shown as sticks in the same color as the backbone. **(E)** Co-affinity purification assessing the interaction between His_6_-TmaT and indicated FLAG-AftD variants overexpressed in *E. coli* cells using pCDFDuet-1 and pET22/42 plasmids, respectively. Eluates were analyzed via SDS-PAGE, visualized using CBB staining and α-His/α-FLAG immunoblots. **(F)** Growth curves of *aftD* cKO strains complemented with WT, or the indicated interaction mutants in 7H9 liquid medium in the absence or presence (inset) of acetamide. **(G)** Representative TLC analysis of total extractable lipids isolated from the *aftD* cKO strain grown in the presence (+) or absence (-) of acetamide to control *aftD* expression, additionally complemented with an empty vector pJEB402 (EV) or plasmids constitutively expressing WT AftD, or the indicated active site/interaction variants. **(H)** TMM:TDM ratios were quantified from the TLCs in (E) and averaged across three independent biological replicates, and shown with error bars (± SD). **(I)** Representative TLC analyses of [^14^C]-labeled sugars released after hydrolysis from the insoluble cell wall-linked mAGP fractions isolated from *aftD* cKO strains grown in the presence (+) or absence (-) of acetamide to control *aftD* expression, additionally complemented with an empty vector pJEB402 (EV) or plasmids constitutively expressing indicated mutants. Right side is the Ara:Glc ratios were quantified and averaged across three independent biological replicates, and shown with error bars (± SD). Student’s *t*-test: ns, not significant; **, *P* < 0.01; ****, *P* < 0.0001. Where a target gene depleted, red (“-” acetamide) symbols are used.

### The physical TmaT-AftD interaction is crucial for its function in ensuring proper mAGP assembly

We also performed targeted mutagenesis to experimentally validate the TmaT-AftD interface, systematically changing 14 polar residues on or close to TM11’ of AftD, and within the interfacial helix at the base of TM 11’ (6 Å of TmaT), to non-polar amino acids (Figure 6D and S9A). All these AftD substitution variants could be robustly co-purified with TmaT in *E. coli*, except AftD^D386A^, where the TmaT-AftD interaction was substantially impaired (Figure S9A and 6E). As this alanine substitution specifically disrupted binding without affecting protein stability (Figure S9), we conclude that the Asp386 is an essential determinant of TmaT-AftD complex formation; Asp386 is located in the linker between TM11’ and the interfacial helix (Figure 6D), possibly influencing how this pair of helices is presented for interaction with TmaT, perhaps via salt bridges with one of the adjacent positively charged residues (i.e. R385) in this linker.

We then assessed the physiological consequences of this interaction-deficient AftD^D386A^ variant in *M. smegmatis*, along with a double substitution variant AftD^R385A/D386A^ that we also constructed. Both variants could be stably produced in cells, similar to WT (Figure S9B). Expression of both *aftD^D386A^* and *aftD^R385A/D386A^* failed to complement the growth of the *aftD* cKO strain when *aftD^WT^*expression was not induced (Figure 6F). Furthermore, the *aftD^R385A/D386A^* double mutant caused a growth defect even when *aftD^WT^* was present (Figure 6F). This loss of cellular function correlated directly with impaired mAGP assembly, as judged by defective TMM transport (accumulated TMM and reduced TDM; Figure 6G and 6H), and ineffective AG incorporation onto the cell wall (reduced arabinose and galactose in mAGP; Figure 6I). Together with our catalytic site mutagenesis data, these results establish that a functional, physically-coupled TmaT-AftD complex is indispensable for coordinating TMM transport and AG ligation to furnish the mAGP architecture.

## Discussion

The biosynthesis of the mycobacterial mAGP complex is essential for cell viability, and requires tight coordination of AG synthesis and MA transport. Yet, how the cell synchronizes these two pathways has been unexplored. In this study, we have discovered a direct interaction between the AG arabinofuranosyltransferase AftD and the putative TMM acyltransferase TmaT, providing direct evidence for this integration. We have established that AftD is required for regulating TMM transport, in a manner dependent on both its catalytic activity and its interaction with TmaT. We have further demonstrated that both AftD and TmaT are required for AG biosynthesis in the cell envelope, specifically ligation to the peptidoglycan layer. The TmaT-AftD complex defines a novel regulatory node that coordinates AG synthesis and MA transport pathways, enabling proper mycobacterial cell envelope assembly. Of note, a parallel study showed that the same complex is conserved in *Corynebacterium glutamicum*, suggesting a common and/or related function across the *Mycobacteriales* order^33^.

Our discovery reveals previously unappreciated insights into the possible role of TmaT in mycobacteria, especially as part of the TmaT-AftD regulatory node. It has been proposed that TmaT acetylates TMM in the cytoplasmic leaflet of the IM to produce the Ac-TMM substrate for transport across the membrane by MmpL3 homologs in *Corynebacteria*^31^. However, this is inconsistent with our structural evidence, where acetyl-CoA is bound in TmaT in a manner that points the acetyl group towards the periplasmic leaflet, suggesting a periplasmic substrate. It is possible that TmaT generates periplasmic Ac-TMM instead, though it is not obvious how TMM can access the active site from within the membrane. If indeed formed, regardless, periplasmic Ac-TMM may be required for MmpL3-mediated release from the IM. Here, AftD may provide a structural scaffold for TmaT, stabilizing its fold and/or activating enzymatic function. It is additionally possible that given their proximity, AftD enzymatic substrates or products may inhibit or activate TmaT function, respectively, ultimately impacting TMM transport. Taking a completely different angle, TMM may not actually be the substrate for TmaT in mycobacteria. Other AT3 family enzymes utilize cytoplasmic acyl-CoA to acylate diverse extracytoplasmic carbohydrates^46^, including peptidoglycan (OatA)^45,47^, O-antigen (OafB)^41^ and the enterobacterial common antigen (WecH)^48^. Given its interaction with AftD, it is conceivable that AG biosynthetic intermediates may instead be possible substrates for TmaT acetylation, and such a modification could be required for the transfer of AG polysaccharides onto peptidoglycan. The lack of mature AGP architecture may then subsequently affect TMM transport. In fact, how AftD itself is involved in AG biosynthesis still require clarification^17,18^; yet heretofore unknown, we have demonstrated that its activity also directly impacts the ligation of AG onto the cell wall. Further investigation will be necessary to fully elucidate how the TmaT-AftD complex regulates mAGP assembly in mycobacteria.

The TmaT-AftD complex acts as a molecular node for synchronizing cell envelope assembly, specifically coordinating MA transport and AG ligation to the cell wall. This physical coupling ensures the proper maturation of the mAGP complex, thus the mycobacterial OM, reminiscent of regulatory strategies seen in *Pseudomonas aeruginosa*, where direct interaction between cell wall and lipopolysaccharide biosynthetic enzymes coordinates assembly of the Gram-negative bacterial OM^49^. The TmaT-AftD complex therefore represents a novel, attractive target for future anti-mycobacterial therapeutics.

## Materials and Methods

### Bacterial strains and growth conditions

All bacterial strains used in this study are listed in Table S3. *Mycobacterium smegmatis* mc²155 was used as the parental strain for all experiments. The conditional knockout strains for *tmaT* (*MSMEG_0319*) and *aftD* (*MSMEG_0359*) were derived from the WT mc²155 genetic background. *M. smegmatis* was cultured at 37 °C in Middlebrook 7H9 broth (BD Difco, 271310) supplemented with 0.5% glycerol and 0.05% tyloxapol (Merck, T0307), or in Tryptic Soy Broth (TSB; Merck, 1.05459) containing 0.05% Tween 80 (Merck, P8074). Solid cultures were grown on LB agar (Merck, L3022). Antibiotics for *M. smegmatis* were used at the following concentrations: 25 μg/mL kanamycin (Kan; Sigma-Aldrich), 20 μg/mL spectinomycin (Spec; Sigma-Aldrich), and 50 μg/mL hygromycin (Hyg; Merck).

*E. coli* NovaBlue and BL21(λDE3) (Novagen) were utilized for molecular cloning and protein expression, respectively. *E. coli* strains were grown in LB broth or on LB agar at 37 °C. Antibiotics for *E. coli* selection were: 50 μg/mL Kan, 50 μg/mL Spec, 150 μg/mL Hyg, and 100 μg/mL ampicillin (Amp; Sigma-Aldrich).

### Plasmids and cloning procedures

All plasmids used in this study are detailed in Table S4. DNA fragments were amplified by PCR using KOD One™ PCR Master Mix (Merck, KMM-101), and cloning was performed via restriction digestion with appropriate enzymes (New England Biolabs) followed by ligation using T4 DNA Ligase (New England Biolabs, M0202). Alternatively, constructs were assembled using the Gibson Assembly Master Mix (New England Biolabs, E2611). Plasmids were transformed into *E. coli* NovaBlue competent cells for propagation and purified using the QIAprep Spin Miniprep Kit (Qiagen, 27104). The sequences of all DNA constructs were verified by Sanger sequencing (Bio Basic Asia Pacific, Singapore). Primers used in conditional knock out and mutations are listed in Table S5.

### Generation of *tmaT* and *aftD* conditional knockout, CRISPRi knock down, as well as derivative strains

Conditional knockout (cKO) strains of *M. smegmatis* for *tmaT* and *aftD* were generated via two-step homologous recombination as previously described^50^. Briefly, pYUB854-derived suicide plasmids were constructed harbouring a *spec^R^* cassette flanked by the 5′-and 3′-flanking regions of the target gene (*tmaT* or *aftD*), alongside *sacB* and *lacZ* markers. The suicide plasmids were electroporated (2200 V; Eporator, Eppendorf) into WT *M. smegmatis* mc²155. First-crossover integrants (blue colonies) were selected on LB agar supplemented with 50 µg/mL hygromycin (Hyg; Merck), 20 µg/mL spectinomycin (Spec; Sigma-Aldrich), and 20 µg/mL X-gal (Bio Basic Asia Pacific).

To facilitate the second crossover of these essential genes, a kanamycin-resistant episomal plasmid carrying the target gene under an acetamide-inducible promoter (pMyC-*tmaT* or pMyC-*aftD*) was introduced into the first-crossover cells by electroporation. Transformants were then plated on LB agar containing 5% (w/v) sucrose, 25 µg/mL kanamycin (Kan), 20 µg/mL Spec, and 0.05% (w/v) acetamide (Sigma-Aldrich) to induce the second-crossover event. Potential cKO mutants (white colonies) were screened by colony PCR and the specific in-frame deletions were confirmed by Sanger sequencing (Bio Basic Asia Pacific, Singapore). For complementation and mutant investigation, WT, or mutated variants of *tmaT* or *aftD* were cloned into the integrative plasmid pJEB402-*hyg^R^* under the control of the constitutive mycobacterial optimized promoter (MOP).

### Native affinity purification from *M. smegmatis*

*M. smegmatis* mc²155 cells harboring pJEB402-*his_6_*-*tmaT*, pJEB402-*his_6_*-*aftD* or empty vector (control) were grown in 1.5-L TSB (Merck, 1.05459) supplemented with 0.05% Tween 80 (Merck, P8074) to the stationary phase. Cells were harvested by centrifugation (4,700 *g*, 20 min) and resuspended in 20 mL cold TBS (20 mM Tris-HCl pH 8.0, 150 mM NaCl) containing 100 µg/mL lysozyme (Merck, L6876), 100 µM phenylmethylsulfonyl fluoride, and 50 µg/mL DNase I. After lysis via three passages through a French Press (GlenMills) at 18,000 psi, the suspension was centrifuged at 4,700 *g* for 5 min. The resulting supernatant was subjected to ultracentrifugation at 95,000 x *g* for 1 hour. The pelleted membrane was resuspended in 10 mL extraction buffer (TBS pH 8.0 containing 5 mM MgCl_2_, 10% glycerol, 5 mM imidazole, and 1% n-dodecyl-β-D-maltoside (DDM; Calbiochem)) and extracted on ice for 2 h. The solubilized fraction was clarified by ultracentrifugation at 95,000 x *g* for 1 hour and loaded onto 500 µL TALON metal affinity resin (Takara Bio, 635504). The resin was washed ten times with 500 µL Wash Buffer (20 mM Tris-HCl pH 8.0, 300 mM NaCl, 5 mM MgCl_2_, 10% glycerol, 8 mM imidazole, and 0.05% DDM). Bound proteins were eluted in four 100 µL fractions using elution buffer (TBS pH 8.0, 5 mM MgCl_2_, 10% glycerol, 200 mM imidazole, and 0.05% DDM). Eluates were concentrated using an Amicon Ultra 30 kDa centrifugal filter (Merck, UFC501096) and resolved by SDS-PAGE. Protein bands were visualized by staining with Coomassie Brilliant Blue (Merck, 1.12553), excised and identified by LC-MS/MS (Taplin Biological Mass Spectrometry Facility, Harvard Medical School).

### Heterologous co-expression and pull-down from *E. coli*

*E. coli* BL21(λDE3) cells were co-transformed with pCDF plasmids expressing His-tagged TmaT (wild-type or mutants) and pET22/42 plasmids expressing FLAG-tagged AftD (wild-type or mutants). Cultures were grown in LB medium at 37 °C to an OD_600_ of 0.6–0.8, induced with 1 mM isopropyl β-d-thiogalactopyranoside (IPTG; Axil Scientific), and the cell culture was grown for another 3hr. Harvested cells will be subjected to the similar processes for affinity co-purification from *M. smegmatis*.

### Growth curve measurements

*M. smegmatis* mc²155 *tmaT* or *aftD* cKO strains, with or without either WT or mutant complementation plasmids, were pre-cultured in Middlebrook 7H9 broth (BD Difco, 271310) supplemented with 0.5% glycerol and 0.05% tyloxapol (Merck, T0307), and 0.05% (w/v) acetamide (Sigma-Aldrich, A0500) at 37 °C, respectively. Upon reaching mid-log phase, cells were harvested and washed twice with 7H9 broth containing 0.05% tyloxapol to remove residual acetamide. Cultures were then back-diluted to an initial OD_600_ of 0.001 in 10 mL of 7H9 broth. Growth was monitored in the presence or absence of acetamide (to induce gene depletion) by measuring the OD_600_ three times daily over a 72-hour period. For CRISPRi-mediated knockdown, *M. smegmatis* WT and derivative strains harboring CRISPRi constructs were pre-cultured in 7H9 broth supplemented with 0.5% glycerol and 0.05% tyloxapol. Upon reaching mid-exponential phase, cells were back-diluted to an initial OD_600_ of 0.001 in 10 mL of 7H9 broth. To initiate gene knockdown, cultures were grown in the presence of 50 ng/mL or 100ng/ml (only for *tmaT*) anhydrotetracycline (ATc; Sigma-Aldrich, 37919) to induce the expression of a catalytically inactive Cas9 (dCas9)^51^. Transcriptional gene silencing was achieved by the dCas9–sgRNA complex, which sterically hinders RNA polymerase during transcriptional initiation or elongation. Growth was monitored by measuring the measuring the OD_600_ three times daily over a 72-hour period.

### Radiolabeling and TLC analysis of mycobacterial lipids

For lipid profiling, *M. smegmatis* cKO or CRISPRi strains were inoculated at an initial OD_600_ of 0.05 and cultured in Middlebrook 7H9 broth at 37 °C under permissive or restrictive conditions (±0.05% acetamide or ±50–100 ng/mL ATc, respectively). Upon reaching early stationary phase, 1-mL of culture was normalized to an OD_600_ of 1.0 and metabolically labeled with [^14^C]-acetate (PerkinElmer, NEC084H001MC; 0.5 µCi/mL) for 2 h at 37 °C. Labeled cells were harvested (5,000 *g*, 10 min) and total extractable lipids were isolated by resuspending the pellet in chloroform: methanol (2:1, v/v). Following three cycles of sonication (30 s, each), a biphasic system was generated by adding methanol and ultrapure water to a final ratio of 1:1:0.8 (v/v/v). The organic phase was collected and dried. To analyze covalently-linked mycolic acids (MAs), the delipidated cell debris was dried using a SpeedVac Concentrator Plus (Eppendorf) and subjected to saponification in 40% (w/v) tetrabutylammonium hydroxide (TBAH; Sigma-Aldrich, 178780) at 95 °C for 2 h. Released MAs were acidified with HCl, extracted into n-hexane, and dried^52^. Both extractable lipids and liberated MAs were resuspended in chloroform:methanol (2:1, v/v), spotted onto TLC Silica Gel 60 F_254_ plates (Merck, 1.05554), and developed in chloroform: methanol: water (30:8:1, v/v/v). Radiolabeled species were visualized and quantified by phosphor imaging using a Typhoon FLA 9500 system (GE Healthcare).

### Purification of TmaT-AftD for cryo-EM

*M. smegmatis* cells harboring pJEB402-*his_6_*-*tmaT* were grown in 20 mL tryptic soy broth (TSB; Merck, 1.05459) supplemented with 0.05% Tween 80 (Merck, P8074) to the stationary phase (OD_600_ ∼5). This seed culture was inoculated into 4 flasks of 1.5 L TSB. The cells were grown again for 1.5 days to the stationary phase and harvested by centrifugation at 8,000 x *g* for 10 min. Harvested cells were resuspended in cold 100 mL TBS (Tris-buffered saline, 20 mM Tris-HCl pH 8.0, 150 mM NaCl) containing 100 μg/mL lysozyme (Merck, L6876), 100 µM phenylmethylsulfonyl fluoride, and 50 µg/mL DNase I, and lysed by two passages through a French Press (GlenMills) at 18,000 psi. The resulting suspension was centrifuged at 8,000 x *g* for 5 min to remove unbroken cells. The supernatant containing the cell lysate was subjected to centrifugation at 160,000 x *g* for 30 min. The pelleted membrane was resuspended in 35 mL extraction buffer (TBS pH 8.0 containing 5 mM MgCl_2_, 10% glycerol, 1 mM imidazole, and 1% lauryl maltose neopentyl glycol (LMNG; Anatrace, NG310) and extracted on ice with gentle shaking for 1.5 h. The suspension was subjected to a second round of centrifugation at 160,000 x *g* for 30 min. The supernatant was loaded onto 150 μL TALON metal affinity resin (Takara Bio, 635504) pre-equilibrated with TBS pH 8.0 containing 5 mM imidazole. The mixture was allowed to drain by gravity and the filtrate was loaded back to the column and drained again. The resin was washed four times each with 600 μL Washing Buffer A (20 mM Tris-HCl pH 8.0, 300 mM NaCl, 5% glycerol, 2 mM imidazole). The resin was then incubated for 10 min with 4 mg peptidisc in 4 mL 20 mM Tris-HCl pH 8.0. The mixture was drained by gravity again. The resin was washed four times each with 600 μL Washing Buffer B (20 mM Tris-HCl pH 8.0, 300 mM NaCl, 2 mM imidazole). TmaT-AftD in peptidiscs was eluted three times each with 150 μL elution buffer (TBS pH 8.0 containing 200 mM imidazole). The eluate was dialyzed into TBS pH 8.0 to remove imidazole and concentrated to around 30 μL using an Amicon Ultra 100 kDa centrifugal filter (Merck, UFC510096).

### Cryo-EM grid preparation

3 μL freshly prepared TmaT-AftD in peptidiscs (2.9 mg/mL including some non-specifically purified proteins) in the presence or absence of 0.1 mM acetyl-CoA were applied to glow-discharged Quantifoil R1.2/1.3 300 mesh holey carbon grids. The grids were blotted for 5 s at 4 ℃ and 100% relative humidity, and then plunge-frozen in liquid ethane cooled by liquid nitrogen using a Vitrobot system (Thermo Fisher Scientific). The grids were stored in liquid nitrogen until their assembly into autogrids for data acquisition.

### Cryo-EM data acquisition and processing

EM movies were collected on a Titan Krios G4 electron microscope (Thermo Fisher Scientific) operated at 300 kV, equipped with a Falcon 4 direct electron detector and a Selectris energy filter. The data collection was done using an EPU interface with a 20-eV slit width of the energy filter and a nominal magnification of 165,000x. Further details about the data acquisition can be found in Table S6.

The data processing was done in CryoSPARC 4.7^53^. Briefly, the collected movies were pre-processed in CryoSPARC Live which integrates patch motion correction, CTF estimation and particle extraction using blob picking. Some micrographs were rejected based on the relative ice thickness, global motion etc. The extracted particles then went through a few rounds of 2D classfication and selection. Ab initio reconstruction was used on the selected particles to generate the initial volume, which was then refined with heterogenous refinement and non-uniform refinement functions. The processing procedure is detailed in Figures S2 and S4.

### Model building and refinement

An initial model of the *M. smegmatis* TmaT-AftD complex was generated using AlphaFold3^37^. The sharpened EM density map was then fitted into the model in ChimeraX 1.9^54^. The model was manually fitted into the density using Coot 1.1.20^55^, iteratively evaluated using Phenix 1.21.2^56^ and fine-tuned using the ISOLDE plug-in^57^ in ChimeraX, before the final real-space refinement and evaluation using Phenix.

### Isolation of mAGP fraction from *M. smegmatis*

For the analysis of mAGP-bound sugars, *M. smegmatis* cKO or CRISPRi strains were an initial OD_600_ of 0.05 and cultured at 37 °C under permissive or non-permissive conditions. Cells were harvested at mid-log phase (total OD_600_ = 20), washed twice with 7H9 broth containing 0.05% tyloxapol, and resuspended in 10 mL of the same medium. Cultures were metabolically labeled with 0.5 µCi/mL D-[U-^14^C]-glucose (PerkinElmer, NEC042V050UC) for 3 h at 37 °C ^58^. Labeled cells were harvested, washed with saline, and delipidated by resuspension in chloroform: methanol (2:1, v/v) followed by three cycles of sonication and incubation at 50 °C for 2 h. Following centrifugation (5,000 *g*, 10 min), the resulting pellet was boiled in 50% (v/v) ethanol at 95 °C to remove soluble glycans. The cell wall core was further purified by incubation in 2% (w/v) SDS at 95 °C for 1 h. The insoluble mAGP fraction was recovered by centrifugation. See Supplementary Materials and Methods for isolating and analyzing soluble fraction (50% ethanol).

### Monosaccharide Analysis of the mAGP

Purified mAGP was hydrolyzed by acid hydrolysis in 2 M trifluoroacetic acid (TFA; Sigma-Aldrich, 302031) at 120 °C for 3 h. Hydrolysates were dried using a SpeedVac Concentrator Plus (Eppendorf), resuspended in 100ul ultrapure water, and spotted onto TLC Silica Gel 60 F_254_ plates (Merck, 1.05554). Plates were developed in a solvent system of glacial acetic acid: chloroform: water (7:6:1, v/v/v). Radiolabeled species were visualized and quantified by phosphor imaging using a Typhoon FLA 9500 system (GE Healthcare).

### SDS-PAGE and Immunoblotting

Protein samples were mixed with Laemmli sample buffer (Bio-Rad, 1610747) and resolved by (12%) Tris-glycine SDS-PAGE at 150V for 60 minutes. Gels were either stained with Coomassie Protein Stain (Abcam, ab119211) or processed for immunoblotting. For the latter, proteins were transferred to a polyvinylidene fluoride (PVDF) membrane (Merck, IPVH00010) using a Trans-Blot SD Semi-Dry Transfer Cell (Bio-Rad) at 25 V for 30 min. Membranes were blocked for 2 h at room temperature in 1x Casein Blocking Buffer (Sigma-Aldrich, B6429). To detect tagged proteins, membranes were incubated with horseradish peroxidase (HRP)-conjugated primary antibodies for 1 h. His-tagged proteins were detected using a mouse monoclonal α-Penta-His HRP conjugate (Qiagen, 34460; 1:5,000 dilution), and FLAG-tagged proteins were detected with a mouse monoclonal α-FLAG M2-HRP conjugate (Sigma-Aldrich, A8592; 1:5,000 dilution). Following extensive washing in TBS-T (TBS pH 8.0, 0.1% Tween 20), blots were developed using Luminata Forte Western HRP Substrate (Merck, WBLUF0100).

## Acknowledgements

We thank Jeremy Liang for generating the initial *E. coli* expression plasmid for TmaT used in this study. We thank Jeremy Rock (Rockefeller University) for providing the plRL117 plasmid. The cryo-EM data collection was done at the Cryoelectron Microscopy Facility at the Center for Bioimaging Sciences (NUS); we thank the facility manager Dr. Jian Shi for providing training and technical support. We further thank Dr. Alisa Garaeva and Prof. Markus Seeger (University of Zurich) for providing training on single particle cryo-EM data collection and analysis during an institutional exchange for a different project supported by the Singapore Ministry of Health National Medical Research Council under its Open Fund Young Individual Research Grant (MOH-001624) (to Y.C.). R.H.S. was supported by the NUS Integrative Science and Engineering Program-SCELSE scholarship. This project was funded by the Singapore Ministry of Education Academic Research Fund Tier 2 grant MOE-000702 (S.-S.C.).

## Data Availability

3D cryo-EM maps of apo AftD-TmaT and AftD-TmaT with acetyl-CoA bound have been deposited in the Electron Microscopy Data Bank under accession numbers **EMD-68514** and **EMD-68526**, respectively. Two atomic coordinate files have also been deposited in the Protein Data Bank under the accession numbers **22NM** (AftD-TmaT) and **22NS** (AftD-TmaT_acetyl-CoA).

## References

1. WHO. Global Tuberculosis Report (World Health Organization. https://iris.who.int/server/api/core/bitstreams/e97dd6f4-b567-4396-8680-717bac6869a9/content (2025).

2. Johansen, M. D., Herrmann, J.-L. & Kremer, L. Non-tuberculous mycobacteria and the rise of Mycobacterium abscessus. Nat. Rev. Microbiol. 18, 392–407 (2020).

3. Brennan, P. J. & Nikaido, H. The envelope of mycobacteria. Annu. Rev. Biochem. 64, 29–63 (1995).

4. Dulberger, C. L., Rubin, E. J. & Boutte, C. C. The mycobacterial cell envelope — a moving target. Nat. Rev. Microbiol. 18, 47–59 (2020).

5. Abrahams, K. A. & Besra, G. S. Mycobacterial cell wall biosynthesis: a multifaceted antibiotic target. Parasitology 145, 116–133 (2018).

6. Clifton E. Barry, Dean C. Crick, & Michael R. McNeil. Targeting the Formation of the Cell Wall Core of M. tuberculosis. Infect. Disord. - Drug Targets 7, 182–202 (2007).

7. Beláňová, M. et al. Galactosyl Transferases in Mycobacterial Cell Wall Synthesis. J. Bacteriol. 190, 1141–1145 (2008).

8. Alderwick, L. J., Harrison, J., Lloyd, G. S. & Birch, H. L. The Mycobacterial Cell Wall— Peptidoglycan and Arabinogalactan. Cold Spring Harb. Perspect. Med. 5, a021113 (2015).

9. Alderwick, L. J., Seidel, M., Sahm, H., Besra, G. S. & Eggeling, L. Identification of a Novel Arabinofuranosyltransferase (AftA) Involved in Cell Wall Arabinan Biosynthesis in *Mycobacterium tuberculosis*\*. J. Biol. Chem. 281, 15653–15661 (2006).

10. Belanger, A. E. et al. The embAB genes of Mycobacterium avium encode an arabinosyl transferase involved in cell wall arabinan biosynthesis that is the target for the antimycobacterial drug ethambutol. Proc. Natl. Acad. Sci. 93, 11919–11924 (1996).

11. Telenti, A. et al. The emb operon, a gene cluster of Mycobacterium tuberculosis involved in resistance to ethambutol. Nat. Med. 3, 567–570 (1997).

12. Birch, H. L. et al. Biosynthesis of mycobacterial arabinogalactan: identification of a novel α(1→3) arabinofuranosyltransferase. Mol. Microbiol. 69, 1191–1206 (2008).

13. Herrera, N. et al. The phenotypic landscape of the mycobacterial cell. BioRxiv Prepr. Serv. Biol. 2025.11.14.688347 (2026) doi:10.1101/2025.11.14.688347.

14. Seidel, M. et al. Identification of a Novel Arabinofuranosyltransferase AftB Involved in a Terminal Step of Cell Wall Arabinan Biosynthesis in Corynebacterianeae, such as *Corynebacterium glutamicum* and *Mycobacterium tuberculosis*\*. J. Biol. Chem. 282, 14729– 14740 (2007).

15. Klevorn, T. et al. A periplasmic protein complex mediates arabinofuranosyltransferase activity and intrinsic drug resistance in Mycobacterium tuberculosis. Sci. Adv. 12, eaec5100 (2026).

16. Lissner, R. et al. A new role for lipoproteins LpqZ and FecB in orchestrating mycobacterial cell envelope biogenesis. mBio 17, e0211925 (2026).

17. Škovierová, H. et al. AftD, a novel essential arabinofuranosyltransferase from mycobacteria. Glycobiology 19, 1235–1247 (2009).

18. Alderwick, L. J. et al. AftD functions as an α1 → 5 arabinofuranosyltransferase involved in the biosynthesis of the mycobacterial cell wall core. Cell Surf. 1, 2–14 (2018).

19. Harrison, J. et al. Lcp1 Is a Phosphotransferase Responsible for Ligating Arabinogalactan to Peptidoglycan in Mycobacterium tuberculosis. mBio 7, 10.1128/mbio.00972-16 (2016).

20. Belisle, J. T. et al. Role of the Major Antigen of Mycobacterium tuberculosis in Cell Wall Biogenesis. Science 276, 1420–1422 (1997).

21. Takayama, K., Wang, C. & Besra, G. S. Pathway to Synthesis and Processing of Mycolic Acids in Mycobacterium tuberculosis. Clin. Microbiol. Rev. 18, 81–101 (2005).

22. Marrakchi, H., Lanéelle, M.-A. & Daffé, M. Mycolic Acids: Structures, Biosynthesis, and Beyond. Chem. Biol. 21, 67–85 (2014).

23. Portevin, D. et al. A polyketide synthase catalyzes the last condensation step of mycolic acid biosynthesis in mycobacteria and related organisms. Proc. Natl. Acad. Sci. 101, 314–319 (2004).

24. Chen, Y., Lim, J. D. T. & Chng, S.-S. Exquisite specificity of Pks13 defines the essentiality of trehalose in mycobacteria. Proc. Natl. Acad. Sci. 123, e2507896123 (2026).

25. Xu, Z., Meshcheryakov, V. A., Poce, G. & Chng, S.-S. MmpL3 is the flippase for mycolic acids in mycobacteria. Proc. Natl. Acad. Sci. 114, 7993–7998 (2017).

26. Fay, A. et al. Two Accessory Proteins Govern MmpL3 Mycolic Acid Transport in Mycobacteria. mBio 10, 10.1128/mbio.00850-19 (2019).

27. Zhang, B. et al. Crystal Structures of Membrane Transporter MmpL3, an Anti-TB Drug Target. Cell 176, 636–648.e13 (2019).

28. Su, C.-C. et al. Structures of the mycobacterial membrane protein MmpL3 reveal its mechanism of lipid transport. PLoS Biol. 19, e3001370 (2021).

29. Gupta, K. R. et al. An essential periplasmic protein coordinates lipid trafficking and is required for asymmetric polar growth in mycobacteria. eLife 11, e80395 (2022).

30. Liang, J., Chen, Y. & Chng, S.-S. Defining a critical role of an essential membrane protein in mycolic acid transport in mycobacteria. 2023.05.02.539095 Preprint at 10.1101/2023.05.02.539095 (2023).

31. Yamaryo-Botte, Y. et al. Acetylation of Trehalose Mycolates Is Required for Efficient MmpL-Mediated Membrane Transport in Corynebacterineae. ACS Chem. Biol. 10, 734–746 (2015).

32. Belardinelli, J. M. et al. Structure–Function Profile of MmpL3, the Essential Mycolic Acid Transporter from Mycobacterium tuberculosis. ACS Infect. Dis. 2, 702–713 (2016).

33. Anastacia R. Parks, et al. A TmaT-AftD interaction is required for the CmpL4 mycomembrane biogenesis pathway in Corynebacterium glutamicum. bioRxiv (2026).

34. Takayama, K. & Kilburn, J. O. Inhibition of synthesis of arabinogalactan by ethambutol in Mycobacterium smegmatis. Antimicrob. Agents Chemother. 33, 1493–1499 (1989).

35. Takayama, K., Armstrong, E. L., Kunugi, K. A. & Kilburn, J. O. Inhibition by ethambutol of mycolic acid transfer into the cell wall of Mycobacterium smegmatis. Antimicrob. Agents Chemother. 16, 240–242 (1979).

36. Carlson, M. L. et al. Profiling the Escherichia coli membrane protein interactome captured in Peptidisc libraries. eLife 8, e46615 (2019).

37. Abramson, J. et al. Accurate structure prediction of biomolecular interactions with AlphaFold 3. Nature 630, 493–500 (2024).

38. Tan, Y. Z. et al. Cryo-EM Structures and Regulation of Arabinofuranosyltransferase AftD from Mycobacteria. Mol. Cell 78, 683–699.e11 (2020).

39. Ma, D. et al. Crystal structure of a membrane-bound O-acyltransferase. Nature 562, 286–290 (2018).

40. Coupland, C. E., Ansell, T. B., Sansom, M. S. P. & Siebold, C. Rocking the MBOAT: Structural insights into the membrane bound O-acyltransferase family. Curr. Opin. Struct. Biol. 80, 102589 (2023).

41. Newman, K. E. et al. A novel fold for acyltransferase-3 (AT3) proteins provides a framework for transmembrane acyl-group transfer. eLife 12, e81547 (2023).

42. Pearson, C. R. et al. Acetylation of Surface Carbohydrates in Bacterial Pathogens Requires Coordinated Action of a Two-Domain Membrane-Bound Acyltransferase. mBio 11, 10.1128/mbio.01364-20 (2020).

43. Thanweer, F. & Verma, N. K. Identification of critical residues of the serotype modifying O-acetyltransferase of Shigella flexneri. BMC Biochem. 13, 13 (2012).

44. Krissinel, E. & Henrick, K. Inference of macromolecular assemblies from crystalline state. J. Mol. Biol. 372, 774–797 (2007).

45. Jones, C. S., Anderson, A. C. & Clarke, A. J. Mechanism of Staphylococcus aureus peptidoglycan O-acetyltransferase A as an O-acyltransferase. Proc. Natl. Acad. Sci. 118, e2103602118 (2021).

46. Pearson, C., Tindall, S., Potts, J. R., Thomas, G. H. & van der Woude, M. W. Diverse functions for acyltransferase-3 proteins in the modification of bacterial cell surfaces. Microbiology 168, 001146 (2022).

47. Bera, A., Herbert, S., Jakob, A., Vollmer, W. & Götz, F. Why are pathogenic staphylococci so lysozyme resistant? The peptidoglycan O-acetyltransferase OatA is the major determinant for lysozyme resistance of Staphylococcus aureus. Mol. Microbiol. 55, 778–787 (2005).

48. Kajimura, J. et al. O Acetylation of the Enterobacterial Common Antigen Polysaccharide Is Catalyzed by the Product of the yiaH Gene of Escherichia coli K-12. J. Bacteriol. 188, 7542–7550 (2006).

49. Hummels, K. R. et al. Coordination of bacterial cell wall and outer membrane biosynthesis. Nature 615, 300–304 (2023).

50. Parish, T. & Stoker, N. G. Use of a flexible cassette method to generate a double unmarked Mycobacterium tuberculosis tlyA plcABC mutant by gene replacement. Microbiology 146 (Pt 8), 1969–1975 (2000).

51. Bosch, B. et al. Genome-wide gene expression tuning reveals diverse vulnerabilities of *M. tuberculosis*. Cell 184, 4579–4592.e24 (2021).

52. Hsu, F.-F., Soehl, K., Turk, J. & Haas, A. Characterization of mycolic acids from the pathogen Rhodococcus equi by tandem mass spectrometry with electrospray ionization. Anal. Biochem. 409, 112–122 (2011).

53. Punjani, A., Rubinstein, J. L., Fleet, D. J. & Brubaker, M. A. cryoSPARC: algorithms for rapid unsupervised cryo-EM structure determination. Nat. Methods 14, 290–296 (2017).

54. Meng, E. C. et al. UCSF ChimeraX: Tools for structure building and analysis. Protein Sci. Publ. Protein Soc. 32, e4792 (2023).

55. Emsley, P. & Cowtan, K. Coot: model-building tools for molecular graphics. Acta Crystallogr. D Biol. Crystallogr. 60, 2126–2132 (2004).

56. Adams, P. D. et al. PHENIX: a comprehensive Python-based system for macromolecular structure solution. Acta Crystallogr. D Biol. Crystallogr. 66, 213–221 (2010).

57. Croll, T. I. ISOLDE: a physically realistic environment for model building into low-resolution electron-density maps. Acta Crystallogr. Sect. Struct. Biol. 74, 519–530 (2018).

58. Mikusová, K., Slayden, R. A., Besra, G. S. & Brennan, P. J. Biogenesis of the mycobacterial cell wall and the site of action of ethambutol. Antimicrob. Agents Chemother. 39, 2484–2489 (1995).

