## Supplementary Information for "Discovery of a regulatory node that coordinates cell envelope assembly in mycobacteria"

Supplementary Tables 1-6

Supplementary Methods

Supplementary References

### Supplementary Figures

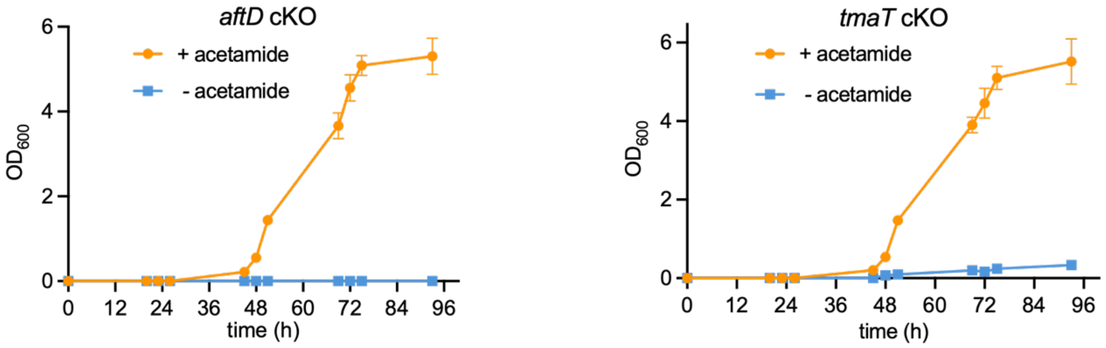


**Figure S1. Depletion of *aftD* or *tmaT* caused growth arrest in *M. smegmatis*.** Growth curves of the *aftD* (*MSMEG_0359*) and *tmaT* (*MSMEG_0319*) conditional knock out (cKO) strains growth curves in 7H9 liquid medium, where gene expression was induced only in the presence or absence of acetamide. Data points represent the mean $OD_{600}$ from three independent biological replicates, with error bars (± SD).

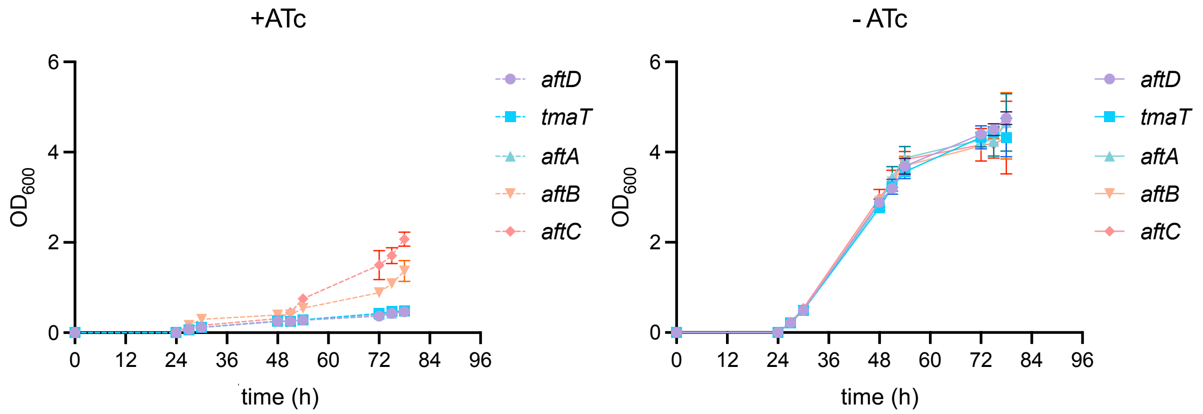


**Figure S2. CRISPRi effectively depletes target genes and impacts growth of *M. smegmatis*.** Growth curves of *M. smegmatis* mc$^2$155 strains upon CRISPRi-mediated knockdown of *aftD*, *tmaT*, *aftA*, *aftB*, or *aftC*. Cultures were grown in the absence (-) or presence (+, to induce gene silencing) of anhydrotetracycline (ATc). Data points represent the mean $OD_{600}$ from three independent biological replicates, with error bars (± SD).

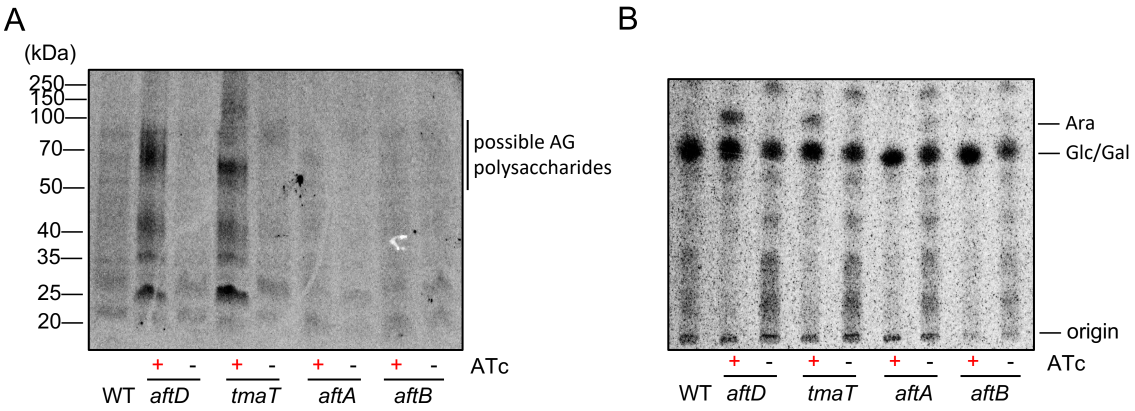


**Figure S3. Depletion of either *aftD* or *tmaT* causes accumulation of putative lipid-linked arabinose-containing polysaccharides. (A)** SDS-PAGE analysis of putative [$^{14}$C]-labeled lipid-linked polysaccharides extracted (50% ethanol) from *M. smegmatis*. Expressiong of the gene is suppressed by CRISPRi when ATc Is present (+). High molecular bands between 50-70 kDa may represent lipid-linked AG before ligation to the cell wall. **(B)** Representative TLC analyses of [$^{14}$C]-labeled sugars (arabinose, Ara; galactose, Gal; glucose, Glc) released after hydrolyzing the 50% ethanol fractions in (A). Where a target gene is depleted, a red ("+" ATc) symbol is used.

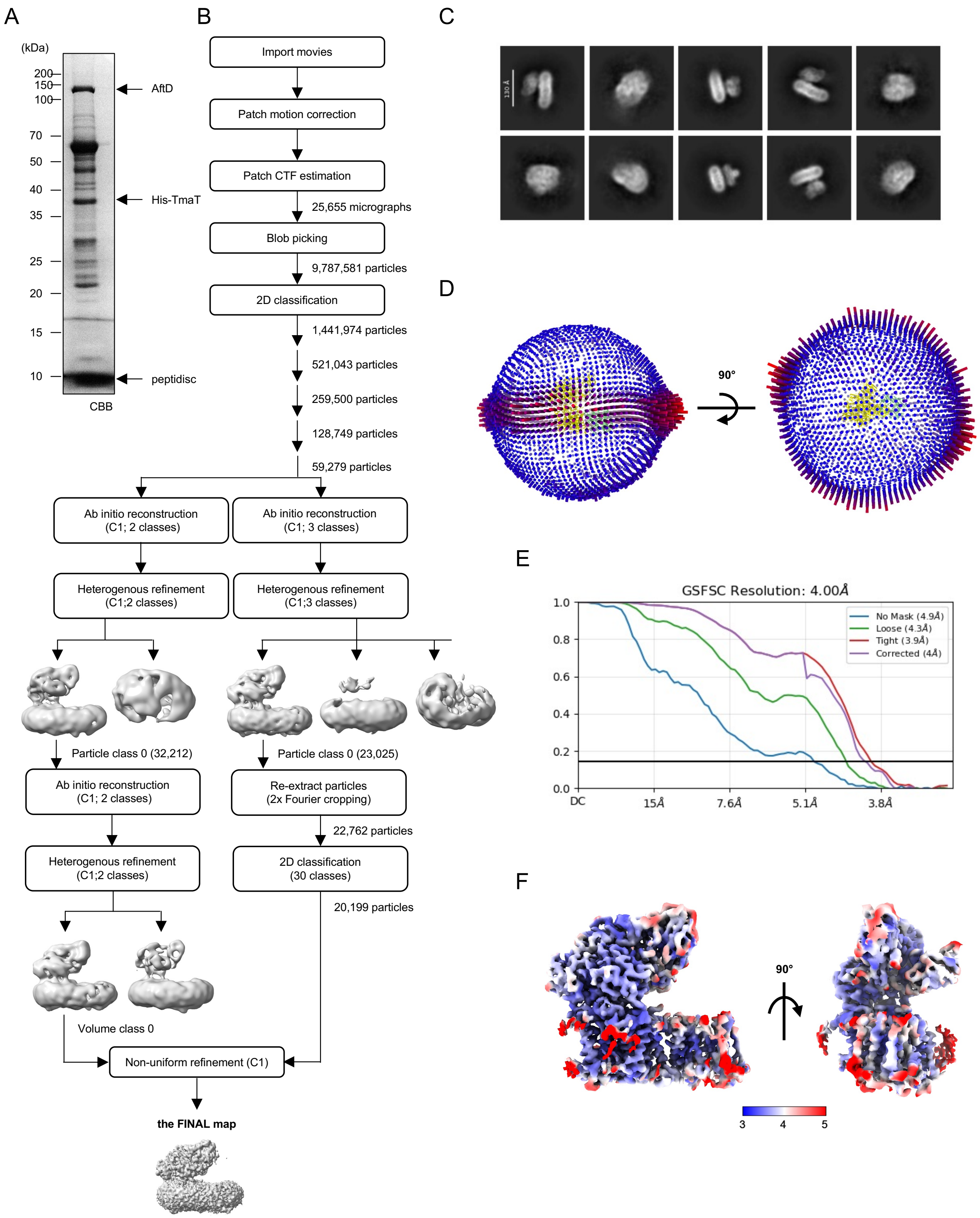


**Figure S4. Single-particle cryo-EM analysis of peptidisc-embedded TmaT-AftD complex. (A)** SDS-PAGE analysis of the TmaT-AftD complex, natively affinity-purified from cells expressing $His_6$-tagged TmaT from an integrative plasmid and reconstituted into peptidiscs. CBB, Coomassie brilliant blue. **(B)** Data processing flowchart yielding the final map (EMD-68514) of the apo TmaT-AftD complex. **(C)** Representative 2D classes derived from 20,199 particles. **(D)** Euler angle distribution plots of the final 3D reconstruction. The height and color of each rod is proportional to the amount of particles visualized from the same specific orientation. **(E)** The gold standard-Fourier Shell Correlation (GS-FSC) plots of unmasked and masked (loose, tight, and corrected) maps derived from cryoSPARC[1]. **(F)** Sharpened local resolution map calculated by cryoSPARC and illustrated with pseudocolor representation of per-voxel resolution.

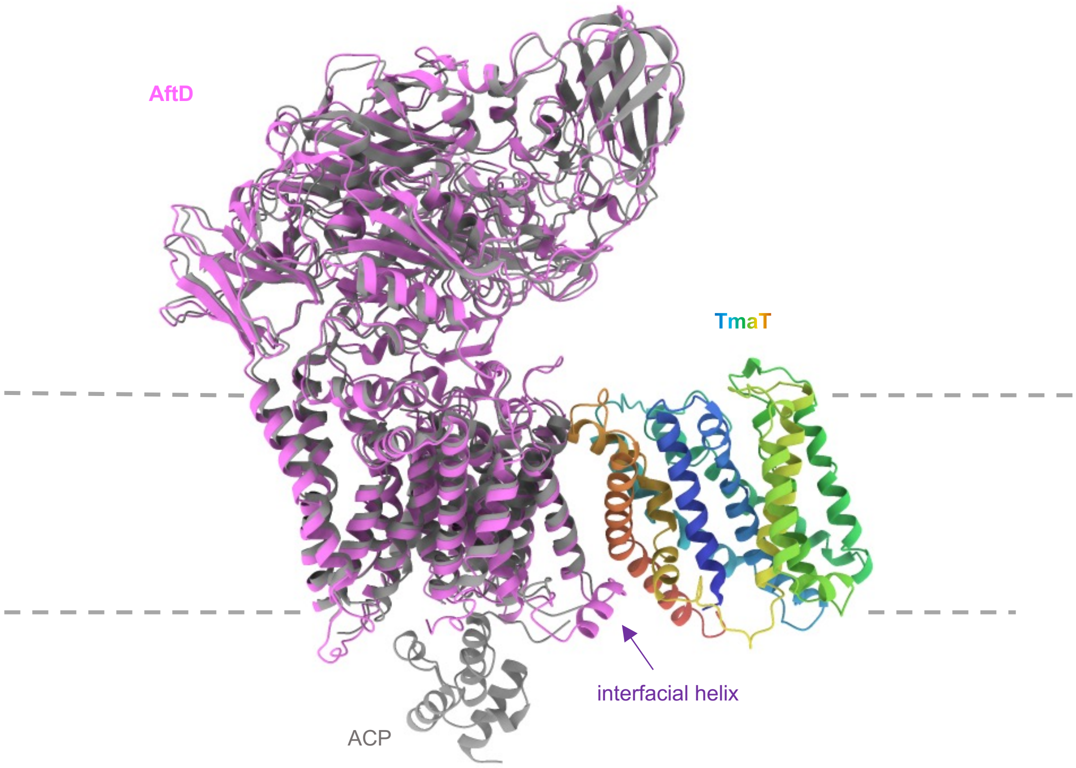


**Figure S5. Structure of the *M. smegmatis* TmaT-AftD complex reveals a conserved AftD architecture devoid of ACP.** Cartoon illustrations of structural overlay of the *M. smegmatis* TmaT-AftD complex (TmaT, rainbow; AftD, pink) and the previously reported *M. abscessus* AftD structure complexed with an acyl carrier protein (ACP) (grey; PDB: 6W98)[2].

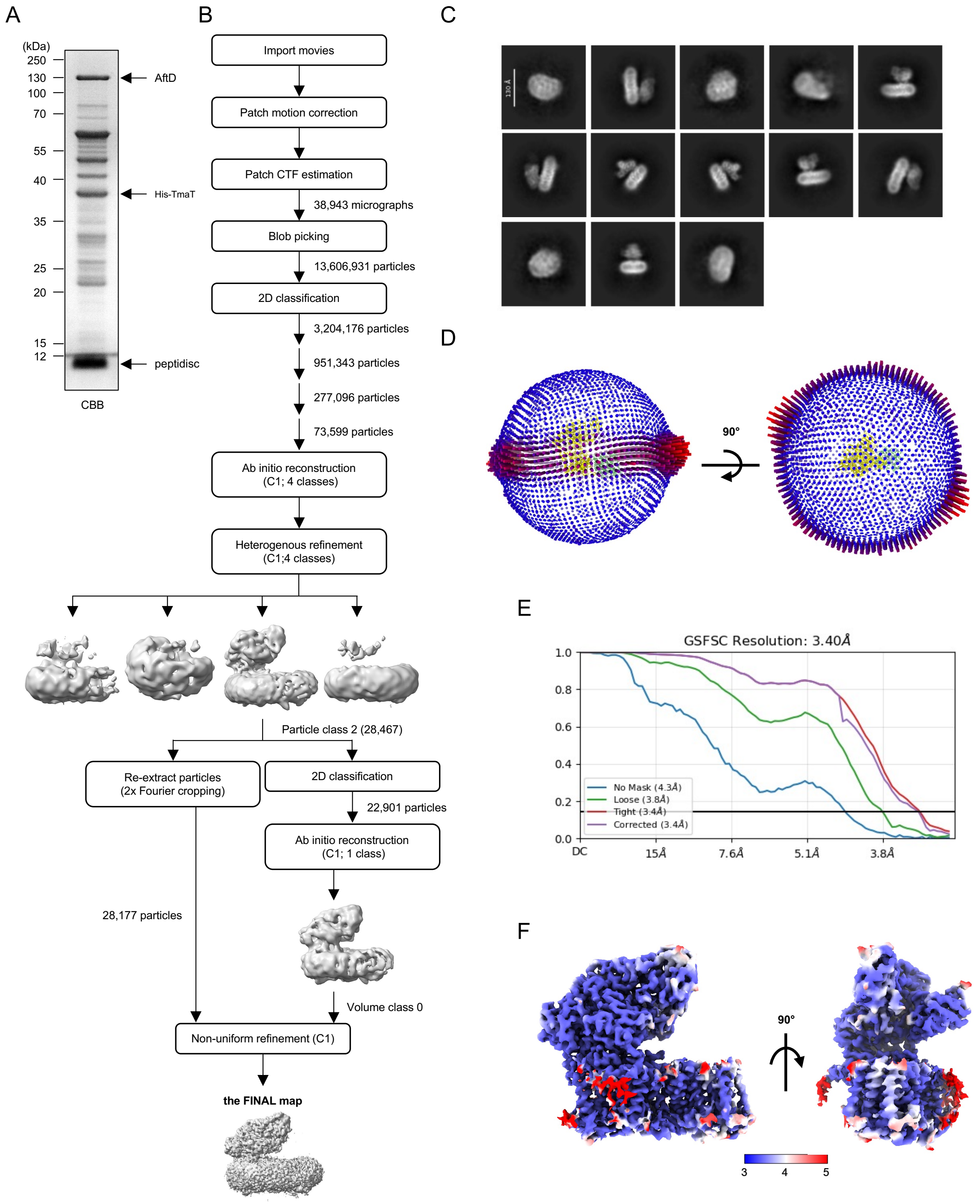


**Figure S6. Single-particle cryo-EM analysis of peptidisc-embedded acetyl-CoA bound TmaT-AftD complex. (A)** SDS-PAGE analysis of the TmaT-AftD complex, natively affinity-purified from cells expressing His$_6$-tagged TmaT from an integrative plasmid and reconstituted into peptidiscs. CBB, Coomassie brilliant blue. **(B)** Data processing flowchart yielding the final map (EMD-68526) of the acetyl-CoA bound TmaT-AftD complex. **(C)** Representative 2D classes derived from 22, 901 particles. **(D)** Euler angle distribution plots of the final 3D reconstruction. The height and color of each rod is proportional to the amount of particles visualized from the same specific orientation. **(E)** The gold standard-Fourier Shell Correlation (GS-FSC) plots of unmasked and masked (loose, tight, and corrected) maps derived from cryoSPARC[1]. **(F)** Sharpened local resolution map calculated by cryoSPARC and illustrated with pseudocolor representation of per-voxel resolution.

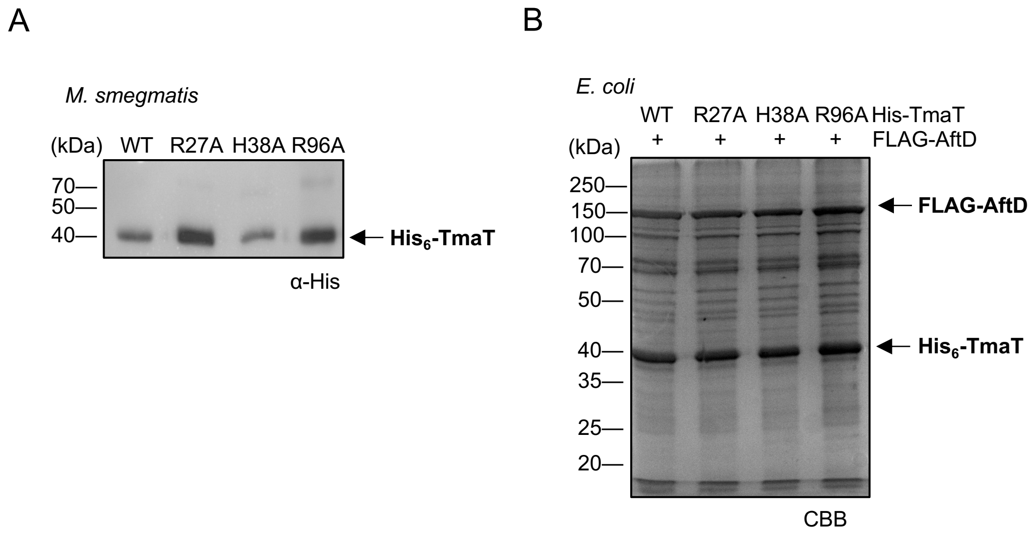


**Figure S7. TmaT variants are stable, and maintain association with AftD. (A)** α-His antibodies immunoblot analysis of total cell lysates of *M. smegmatis* mc²155 cells expressing indicated $His_6$-tagged TmaT variants from the pJEB402 vector. **(B)** Co-affinity purification of indicated $His_6$-TmaT (H-T) variants and FLAG-tagged AftD overexpressed in *E. coli* cells using pCDFDuet-1 and pET22/42 plasmids, respectively. Eluates were analyzed by a SDS-PAGE, visualized using Coomassie brilliant blue (CBB) staining.

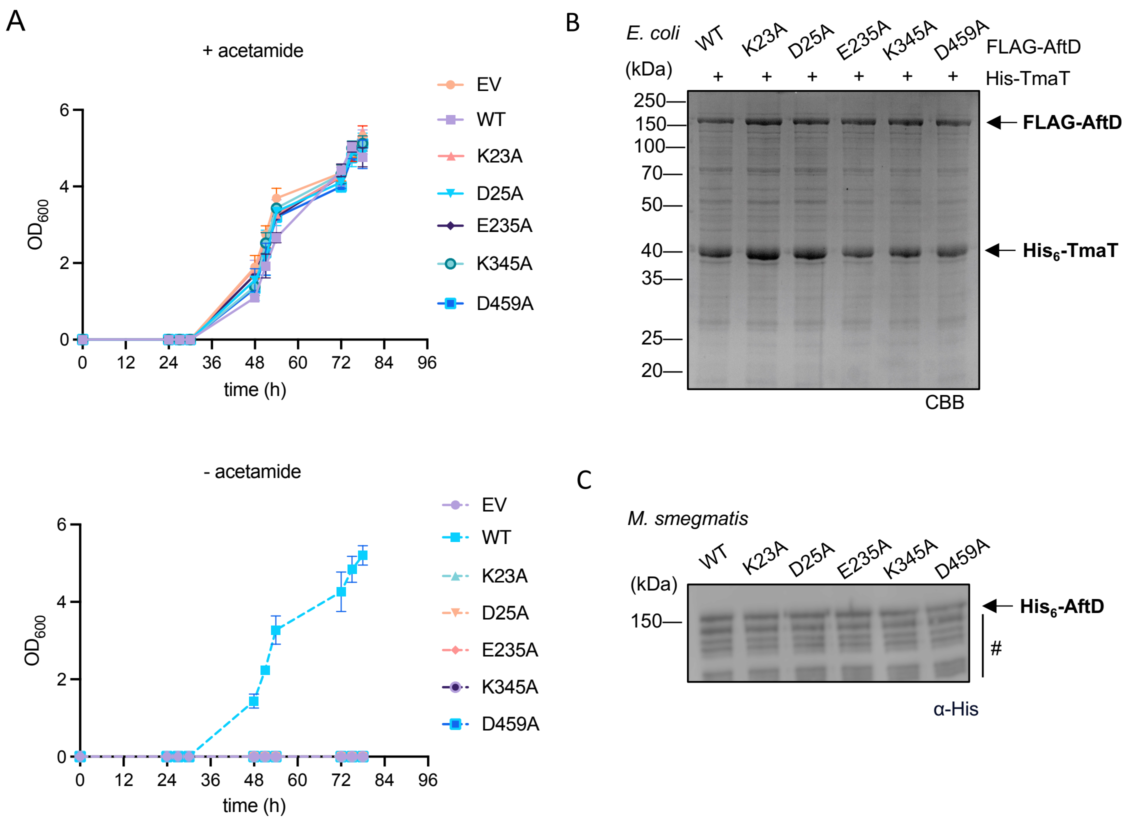


**Figure S8. Predicted active site residues of AftD are essential for function but not for the interaction with TmaT. (A)** Growth curves of the *aftD* cKO strain grown in the presence (+) or absence (-) of acetamide to control *aftD* expression, additionally complemented with an empty vector pJEB402 (EV) or plasmids constitutively expressing WT AftD or the indicated active site mutants. Data represent the mean of three independent biological replicates, with error bars (± SD). **(B)** Co-affinity purification of $His_6$-TmaT and indicated FLAG-tagged AftD variants overexpressed in *E. coli* cells using pCDFDuet-1 and pET22/42 plasmids, respectively. Eluates were analyzed by SDS-PAGE, visualized using CBB staining and α-His/α-FLAG immunoblots. **(C)** Immunoblot analysis of total cell lysates of *M. smegmatis* mc$^2$155 cells expressing indicated $His_6$-tagged AftD variants from the pJEB402 vector, visualized using α-His antibodies. Degradation profiles of AftD variants are similar to WT. Degradation products of AftD are denoted with #.

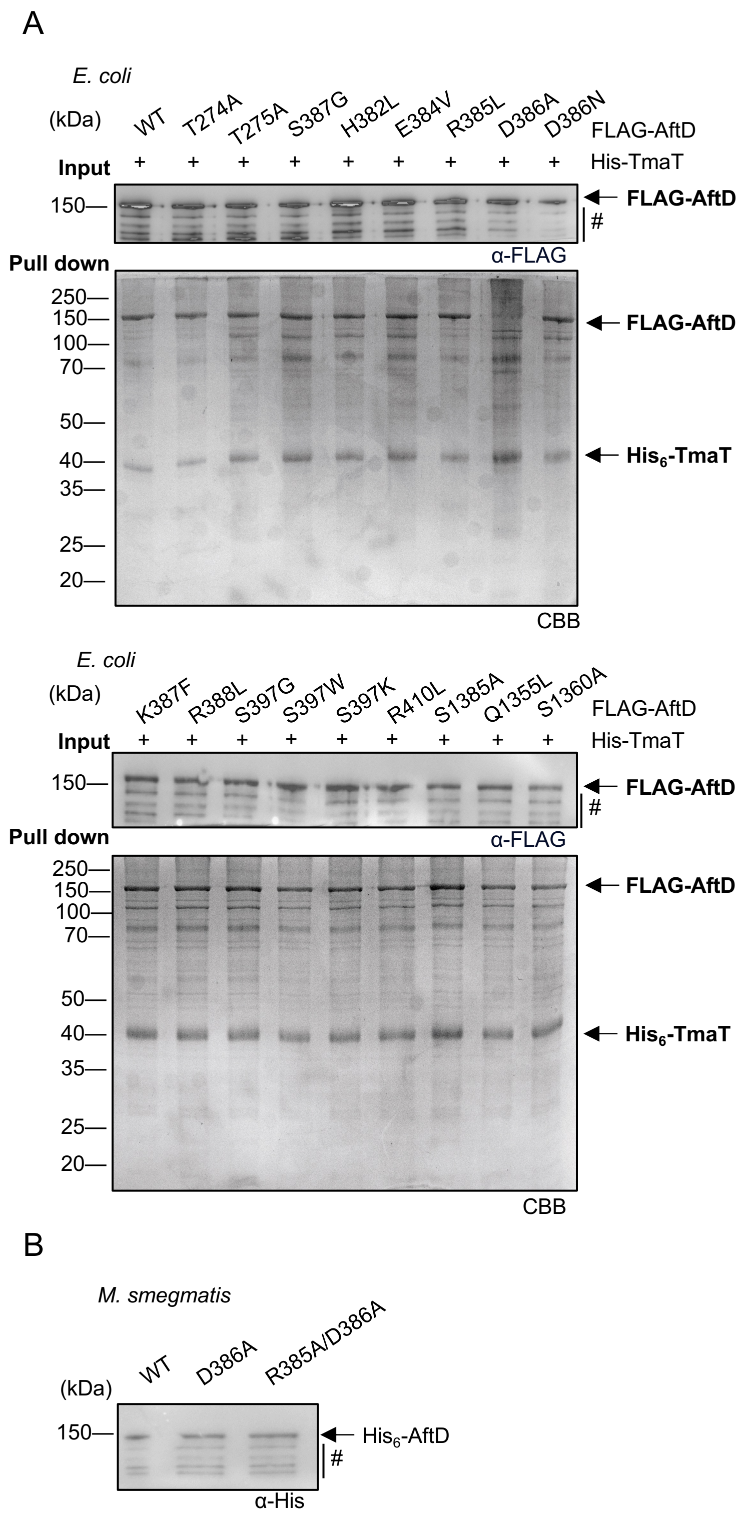


**Figure S9. The D386A mutation on AftD disrupts TmaT-AftD interactions. (A)** SDS-PAGE analyses of proteins affinity purified from *E. coli* cell expressing His$_6$-TmaT and indicated FLAG-tagged AftD variants. The gel was visualized using CBB staining. The top input panel shows the α-FLAG immunoblot analysis of whole cell lysates corresponding to each affinity purified sample. **(B)** α-His immunoblot analysis of total cell lysates of *M. smegmatis* mc²155 cells expressing indicated $His_6$-tagged AftD variants from the pJEB402 vector. Degradation profiles of AftD variants are similar to WT AftD. Degradation products of AftD are denoted with #.

**Supplementary Tables**

**Table S1.** Top protein hits identified in the ~150-kDa band co-purified with His-TmaT in *M. smegmatis*.

| Lane / Protein (kDa) | # peptides detected[a] | |
|---|---|---|
| | pJEB402 | pJEB402-*His-tmaT* |
| **AftD (148)** | **12** | **123** |
| MSMEG_1637 (97) | 3 | 21 |
| Kgd (135) | 4 | 19 |
| LysX (121) | 0 | 18 |

[a] Total number of peptides detected in the excised ~150-kDa gel band, along with corresponding excised region in the control lane. Only proteins with at least 3x more peptides in the sample lane are listed.

**Table S2.** Top protein hits identified in the ~40-kDa band co-purified with His-AftD in *M. smegmatis*.

| Protein (kDa) / Lane | # peptides detected | |
|---|---|---|
| | pJEB402 | pJEB402-*His-aftD* |
| **TmaT (45)** | **0** | **48** |
| GroEL2 (56) | 7 | 25 |
| MSMEG_2612 (39) | 6 | 24 |
| HypB (28) | 4 | 13 |

[a] Total number of peptides detected in the excised ~40-kDa gel band, along with corresponding excised region in the control lane. Only proteins with at least 3x more peptides in the sample lane are listed.

**Table S3.** Bacterial strains used in this study.

| **Strains used** | **Source** |
|---|---|
| *E. coli* BL21(λDE3) | Novagen |
| *E. coli* NovaBlue | Novagen |
| *M. smegmatis* mc$^2$155 | lab collection |
| ***aftD* cKO**: mc$^2$155 Δ*aftD*::*spec*$^R$ pMyC-*aftD* (Kan$^R$) | this study |
| ***tmaT* cKO**: mc$^2$155 Δ*tmaT*::*spec*$^R$ pMyC-*tmaT* (Kan$^R$) | this study |

**Table S4:** Plasmids used in this study.

| Plasmids | Plasmid description | Reference/source |
|---|---|---|
| pJEB402 | Mycobacterial chromosomal integration vector, contains a constitutive MOP promoter and L5 attB site; Kan$^R$ | [3] |
| pJEB402*his*$_6$-*tmaT* | pJEB402; encodes full length MSMEG_0319 with a N- terminal His$_6$ tag; Kan$^R$ | This study |
| pJEB402*his*$_6$-*aftD* | pJEB402; encodes full length MSMEG_0359 with a N- terminal His$_6$ tag; Kan$^R$ | This study |
| pJEB402-*hyg*$^R$ | Derivative of pJEB402; carries a hygromycin resistance cassette in place of the original kanamycin marker; Hyg$^R$ | Lab collection |
| pJEB402-*hyg*$^R$-*aftD* | Derivative of pJEB402; encodes full length *aftD* (MSMEG_0359); Hyg$^R$ | This study |
| pJEB402-*hyg*$^R$-*aftD*$^{K23A}$ | Derivative of pJEB402; encoding full-length *aftD* (MSMEG_0359) with a K23A mutation; Hyg$^R$ | This study |
| pJEB402-*hyg*$^R$-*aftD*$^{D25A}$ | Derivative of pJEB402; encoding full-length *aftD* (MSMEG_0359) with a D25A mutation; Hyg$^R$ | This study |
| pJEB402-*hyg*$^R$-*aftD*$^{E235A}$ | Derivative of pJEB402; encoding full-length *aftD* (MSMEG_0359) with a E235A mutation; Hyg$^R$ | This study |
| pJEB402-*hyg*$^R$-*aftD*$^{K345A}$ | Derivative of pJEB402; encoding full-length *aftD* (MSMEG_0359) with a K345A mutation; Hyg$^R$ | This study |
| pJEB402-*hyg*$^R$-*aftD*$^{D459A}$ | Derivative of pJEB402; encoding full-length *aftD* (MSMEG_0359) with a D459A mutation; Hyg$^R$ | This study |
| pJEB402-*hyg*$^R$-*aftD*$^{R385A}$ | Derivative of pJEB402; encoding full-length *aftD* (MSMEG_0359) with a R385A mutation; Hyg$^R$ | This study |

| | | |
|---|---|---|
| pJEB402-*hyg*$^{R}$-*aftD*$^{D386A}$ | Derivative of pJEB402; encoding full-length *aftD* (MSMEG_0359) with a D386A mutation; Hyg$^{R}$ | This study |
| pJEB402-*hyg*$^{R}$-*aftD*$^{R385AD386A}$ | Derivative of pJEB402; encoding full-length *aftD* (MSMEG_0359) with a R385A/D386A mutation; Hyg$^{R}$ | This study |
| pJEB402-*hyg*$^{R}$-*aftD*$^{E235AR385AD386A}$ | Derivative of pJEB402; encoding full-length *aftD* (MSMEG_0359) with a E235A/R385A/D386A mutation; Hyg$^{R}$ | This study |
| pJEB402-*hyg*$^{R}$-*tmaT* | Derivative of pJEB402; encodes full length *tmaT* (MSMEG_0319); Hyg$^{R}$ | This study |
| pJEB402-*hyg*$^{R}$-*tmaT*$^{R27A}$ | Derivative of pJEB402; encodes full length *tmaT* (MSMEG_0319) with a R27A mutation; Hyg$^{R}$ | This study |
| pJEB402-*hyg*$^{R}$-*tmaT*$^{H38A}$ | Derivative of pJEB402; encodes full length *tmaT* (MSMEG_0319) with a H38A mutation; Hyg$^{R}$ | This study |
| pJEB402-*hyg*$^{R}$-*tmaT*$^{R96A}$ | Derivative of pJEB402; encodes full length *tmaT* (MSMEG_0319) with a R96A mutation; Hyg$^{R}$ | This study |
| pIRL117 | Mycobacterial chromosomal integration vector; Optimized M. smegmatis CRISPRi plasmid; L5 integrating; Tet-On promoter controlled by ATc; expresses dCas9; Kan$^{R}$ | pIRL117 was a gift from Jeremy Rock (Addgene plasmid # 163635; http://n2t.net/addgene:163635; RRID:Addgene_163635) ; [4] |
| pIRL117-*aftAsgRNA* | pIRL117; expresses of an *aftA*-targeting sgRNA; used for CRISPRi-mediated knockdown; Kan$^{R}$ | This study |
| pIRL117-*aftBsgRNA* | pIRL117; expresses of an *aftB*-targeting sgRNA; used for CRISPRi-mediated knockdown; Kan$^{R}$ | This study |
| pIRL117-*aftCsgRNA* | pIRL117; expresses of an *aftC*-targeting sgRNA; used for CRISPRi-mediated knockdown; Kan$^{R}$ | This study |

| | | |
|---|---|---|
| pIRL117-*aftDsgRNA* | pIRL117; expresses of an *aftD*-targeting sgRNA; used for CRISPRi-mediated knockdown; $Kan^R$ | This study |
| pIRL117-*tmaT*sgRNA | pIRL117; expresses of a *tmaT*-targeting sgRNA; used for CRISPRi-mediated knockdown; $Kan^R$ | This study |
| pMYC | Mycobacteria episomal shuttle vector for M. smegmatis; contains an acetamide-inducible promoter and dual origins of replication (oriM/oriE); $Kan^R$ | [5] |
| pMYC-*tmaT* | pMYC; encoding full length *tmaT* (MSMEG_0319) under the control of an acetamide-inducible promoter; $Kan^R$ | This study |
| pMYC-*aftD* | pMYC; encoding full length *aftD* (MSMEG_0359) under the control of an acetamide-inducible promoter; $Kan^R$ | This study |
| pYUB854 | Suicide vector for homologous recombination in Mycobacteria; contains an *E. coli* origin (oriE) and $\lambda$-cos sites; lacks a mycobacterial origin of replication. | [6] |
| pYUB854-5'3'*tmaT*-lacZsacB | pYUB854; expresses specR cassette flanked by ~1000 bases of the 5'- and 3'-untranslated regions of the MSMEG_0319 gene. | This study |
| pYUB854-5'3'*aftD*-lacZsacB | pYUB854; expresses specR cassette flanked by 1000 bases of the 5'- and 3'-untranslated regions of the MSMEG_0359 gene. | This study |
| pET22/42 | *E. coli* expression vector; pET22b(+) derivative harboring the pET42a(+) multiple cloning site (MCS); features an IPTG-inducible T7-lac promoter; $Amp^R$ | Lab collection |

| | | |
|---|---|---|
| pET22/42—*FLAG-aftD* | pET22/42; encoding full length of aftD (MSMEG_0359) with a C-terminal Flag tag; $Amp^R$ | This study |
| pET22/42-*FLAG-aftD*$^{R385A}$ | pET22/42; encoding full-length *aftD* (MSMEG_0359) with a C-terminal Flag tag and a R385A mutation; $Hyg^R$ | This study |
| pET22/42-*FLAG-aftD*$^{D386A}$ | pET22/42; encoding full-length *aftD* (MSMEG_0359) with a C-terminal Flag tag and a D386A mutation; $Hyg^R$ | This study |
| pET22/42-*FLAG aftD*$^{R385AD386A}$ | pET22/42; encoding full-length *aftD* (MSMEG_0359) with a C-terminal Flag tag and a R385A/D386A mutation; $Hyg^R$ | This study |
| pET22/42-*FLAG-aftD*$^{K23A}$ | pET22/42; encoding full-length *aftD* (MSMEG_0359) with a C-terminal Flag tag and a K23A mutation; $Hyg^R$ | This study |
| pET22/42-*FLAG-aftD*$^{D25A}$ | pET22/42; encoding full-length *aftD* (MSMEG_0359) with a C-terminal Flag tag and a D25A mutation; $Hyg^R$ | This study |
| pET22/42-*FLAG-aftD*$^{E235A}$ | pET22/42; encoding full-length *aftD* (MSMEG_0359) with a C-terminal Flag tag and a E235A mutation; $Hyg^R$ | This study |
| pET22/42-*FLAG-aftD*$^{K345A}$ | pET22/42; encoding full-length *aftD* (MSMEG_0359) with a C-terminal Flag tag and a K345A mutation; $Hyg^R$ | This study |
| pET22/42-*FLAG-aftD*$^{D459A}$ | pET22/42; encoding full-length *aftD* (MSMEG_0359) with a C-terminal Flag tag and a D459A mutation; $Hyg^R$ | This study |
| pCDFDuet-1 | *E. coli* expression vector; designed for the co-expression of two genes under independent T7-lac promoters; contains the CloDF13 origin of | This study |

| | | |
|---|---|---|
| | replication (CDF) for compatibility with pET and other Duet-series vectors; $Spec^R$ | |
| pCDF-*His₆*-*tmaT* | pCDFDuet-1, encodes full length *tmaT* (MSMEG_0319) with a N- terminal $His_6$ tag; $Spec^R$ | This study |
| pCDF-*His₆*-*tmaT*$^{R27A}$ | pCDFDuet-1, encodes full length *tmaT* (MSMEG_0319) with a N- terminal $His_6$ tag and a R27A mutation; $Spec^R$ | This study |
| pCDF-*His₆*-*tmaT*$^{H38A}$ | pCDFDuet-1, encodes full length *tmaT* (MSMEG_0319) with a N- terminal $His_6$ tag and a H38A mutation; $Spec^R$ | This study |
| pCDF-*His₆*-*tmaT*$^{R96A}$ | pCDFDuet-1, encodes full length *tmaT* (MSMEG_0319) with a N- terminal $His_6$ tag and a R96A mutation; $Spec^R$ | This study |

**Table S5.** Primers used in conditional knock out and mutations.

| Primer name | Sequence (5' to 3') |
|---|---|
| pYUB854-*tmaT* 5'F | tcatggtaatacgactcactagtggtagccgtgttcttcgc |
| pYUB854-*tmaT* SmR+5'R | aaggtagtcggcaaataaaagcttggcgaccagtgt |
| pYUB854-*tmaT* 5'+SmR F | acactggtcgccaagcttttatttgccgactacctt |
| pYUB854-*tmaT* 3'+SmR R | gctccggctgctcggtgaaagctttttgtttatttt |
| pYUB854-*tmaT* SmR+3' F | aaaataaacaaaaagctttcaccgagcagccggag |
| pYUB854-*tmaT 3'R* | tttcggacggttgctagcggacggtcgggtgc |
| pYUB854-*aftD* 5'F | ggtaatacgactcactagggcccgggaagtgcttga |
| pYUB854-*aftD* SmR+5'R | ccaaggtagtcggcaaataaaagcttccacagccagcgcc |
| pYUB854-*aftD* 5'+SmR F | ggcgctggctgtggaagcttttatttgccgactaccttgg |
| pYUB854-*aftD* 3'+SmR R | gttgaggtggcaaggattggaagctttttgtttatttttctaa |
| pYUB854-*aftD* SmR+3' F | ttagaaaaataaacaaaaagcttccaatccttgccacctcaac |
| pYUB854-*aftD 3'R* | taatatttcggacggttgctagctcggcgatgcgcgcagag |
| pMYC-*tmaT* frag F | atccgaattcgatatcgcatgaccaccgagcaatccgcg |
| pMYC-*tmaT* frag R | gtgcgaagcttggctatcgcgcggctggggcg |
| pMYC-*tmaT* pla F | cgccccagccgcgcgatagccaagcttcgcac |
| pMYC-*tmaT* pla R | cgcggattgctcggtggtcatgcgatatcgaattcggat |
| pMYC-*aftD* frag F | cgaattcgatatcgcatggtcgccgcggcgacactc |
| pMYC-*aftD* frag R | gtgcgaagcttggctaaccggcgtgagcggat |
| pMYC-*aftD* pla F | atccgctcacgccggttagccaagcttcgcac |

| | |
|---|---|
| pMYC-*aftD* pla R | gagtgtcgccgcggcgaccatgcgatatcgaattcg |
| *aftD*$^{K23A}$F | CCGACACGGCACTCGACCTCACG |
| *aftD*$^{K23A}$R | GGTCGAGTGCCGTGTCGGGTGAG |
| *aftD*$^{D25A}$ F | CGAAACTCGCCCTCACGGCGAAC |
| *aftD*$^{D25A}$ R | CCGTGAGGGCGAGTTTCGTGTCG |
| *aftD*$^{E235A}$ F | CTTCATCGCGTCGTCGGGCGT |
| *aftD*$^{E235A}$ R | CCGACGACGCGATGAAGTCGAGGAA |
| *aftD*$^{K345A}$ F | ACCTCGCGGCACTGGAACCGGT |
| *aftD*$^{K345A}$ R | GTTCCAGTGCCGCGAGGTTGCG |
| *aftD*$^{D459A}$ F | CAGCCACGCCGAGCCGCTGCAG |
| *aftD*$^{D459A}$ R | GCGGCTCGGCGTGGCTGTTGCC |
| *aftD*$^{R385A}$ F | ATCCGGAAGCCGACAAGCGGGTC |
| *aftD*$^{R385A}$ R | CCGCTTGTCGGCTTCCGGATGCGCGAACGC |
| *aftD*$^{D386A}$ F | CCGGAACGCGCCAAGCGGGTCGCGGTC |
| *aftD*$^{D386A}$ R | CCCGCTTGGCGCGTTCCGGATGCG |
| *aftD*$^{R385AD386A}$ F | ATCCGGAAGCCGCTAAGCGGGTCGCGGTC |
| *aftD*$^{R385AD386A}$ R | ACCCGCTTAGCGGCTTCCGGATGCGCGAACGC |
| *tmaT*$^{R27A}$ F | AAGGCATGGCCGCATGCGCGGCCA |
| *tmaT*$^{R27A}$ R | CGCGCATGCGGCCATGCCTTCGAC |
| *tmaT*$^{H38A}$ F | GGTGCTCACCGCCGTCGCGTTCCAG |
| *tmaT*$^{H38A}$ R | ACGCGACGGCGGTGAGCACCAC |
| *tmaT*$^{R96A}$ F | TGCGCTCCGCGATCGTGCGCATCATGCC |
| *tmaT*$^{R96A}$ R | GCGCACGATCGCGGAGCGCAGGTAGTGC |

**Table S6.** Cryo-EM data collection and modeling statistics.

| | **TmaT-AftD**<br>PDB: 22NM<br>EMD-68514 | **TmaT-AftD with acetyl-CoA**<br>PDB: 22NS<br>EMD-68526 |
|---|---|---|
| **Data collection and processing** | | |
| Magnification | 165,000x | |
| Voltage (kV) | 300 | |
| Set defocus range (μm) | -0.8–2.2 | |
| Energy filter slit width (eV) | 20 | |
| Set total dose (e-/Å$^2$) | 50.5 | |
| Pixel size (Å) | 0.76 | |
| Number of micrographs | 25,655 | 38,943 |
| Number of initial particles | 9,787,581 | 13,606,931 |
| Number of particles in the final map | 20,199 | 28,177 |
| Symmetry | C1 | C1 |
| Global resolution (Å) | 4.00 | 3.40 |
| Local Resolution Range (Å) | 3.4–5.4 | 3.4–4.4 |
| Map sharpening b-factor (Å) | -57.82 | -53.01 |
| **Model statistics** | | |
| Map CC | 0.77 | 0.77 |
| Bond length RMSD (Å) | 0.004 | 0.004 |
| Bond angle RMSD (º) | 0.741 | 0.739 |
| MolProbity score | 1.18 | 0.94 |
| Clash score | 1.51 | 0.60 |

| | | |
|---|---|---|
| Rotamer outliers (%) | 0.22 | 0.00 |
| C-β outliers (%) | 0 | 0 |
| Ramachandran plot | | |
| Favored (%) | 95.80 | 96.42 |
| Allowed (%) | 4.20 | 3.58 |
| Outliers (%) | 0.00 | 0.00 |

### Supplementary Methods

#### Expression test in *M. smegmatis*

*M. smegmatis* mc$^2$155 tmaT or aftD cKO strains with either WT or mutant complementation plasmids, were cultured in Tryptic Soy Broth (TSB; Merck, 1.05459) containing 0.05% Tween 80 (Merck, P8074) supplemented with 0.05% tyloxapol (Merck, T0307), and 0.05% (w/v) acetamide (Sigma-Aldrich, A0500) at 37 °C until $OD_{600}$ reach ~4. Cells were harvested by centrifugation at 4,700 g for 10 min. The cell pellet was mixed with Laemmli sample buffer (Bio-Rad, 1610747) and sonicate for 10 min. The sonicated cell pellete was analyzed by SDS-PAGE and analyzed by immunoblotting using α-His (Merck, H1029) antibodies to detect expression.

#### Monosaccharide analysis of 50% ethanol fraction

Dried 50% ethanol fraction was hydrolyzed by acid hydrolysis in 500ul 2 M trifluoroacetic acid (TFA; Sigma-Aldrich, 302031) at 120 °C for 3 h. Hydrolysates were dried using a SpeedVac Concentrator Plus (Eppendorf), resuspended in 50ul ultrapure water, and spotted onto TLC Silica Gel 60 $F_{254}$ plates (Merck, 1.05554). Plates were developed in a solvent system of glacial acetic acid: chloroform: water (7:6:1, v/v/v). Radiolabeled species were visualized and quantified by phosphor imaging using a Typhoon FLA 9500 system (GE Healthcare).

#### SDS-PAGE of radiolabeled 50% ethanol fraction

To analyze soluble glycans, one-third of the dried 50% (v/v) ethanol fraction was resuspended in 20 µL of ultrapure water and subjected to proteolytic digestion using Proteinase K (Merck, P2308; 1 mg/mL). Samples were then mixed with 2x nonreducing Laemmli sample buffer (Bio-Rad, 1610737) and the 20 µL from each samples was resolved by SDS-PAGE using a 12% Tris-glycine gel. Following electrophoresis, the gel was stained with InstantBlue Coomassie Protein Stain (Abcam, ab119211) and dried using the DryEase Mini-Gel Drying System (Invitrogen, NI2020). The dried gel was exposed to a storage phosphor screen for 72 h, and radiolabeled species were visualized and quantified by phosphor imaging using a Typhoon FLA 9500 system (GE Healthcare).

### Supplementary References


1. Punjani, A., Rubinstein, J. L., Fleet, D. J. & Brubaker, M. A. cryoSPARC: algorithms for rapid unsupervised cryo-EM structure determination. *Nat. Methods* **14**, 290–296 (2017).
2. Tan, Y. Z. *et al.* Cryo-EM Structures and Regulation of Arabinofuranosyltransferase AftD from Mycobacteria. *Mol. Cell* **78**, 683-699.e11 (2020).
3. Lee, M. H., Pascopella, L., Jacobs, W. R. & Hatfull, G. F. Site-specific integration of mycobacteriophage L5: integration-proficient vectors for Mycobacterium smegmatis, Mycobacterium tuberculosis, and bacille Calmette-Guérin. *Proc. Natl. Acad. Sci. U. S. A.* **88**, 3111–3115 (1991).
4. Bosch, B. *et al.* Genome-wide gene expression tuning reveals diverse vulnerabilities of *M. tuberculosis*. *Cell* **184**, 4579-4592.e24 (2021).
5. Beckham, K. S. H., Staack, S., Wilmanns, M. & Parret, A. H. A. The pMy vector series: A versatile cloning platform for the recombinant production of mycobacterial proteins in Mycobacterium smegmatis. *Protein Sci. Publ. Protein Soc.* **29**, 2528–2537 (2020).
6. Bardarov, S. *et al.* Specialized transduction: an efficient method for generating marked and unmarked targeted gene disruptions in Mycobacterium tuberculosis, M. bovis BCG and M. smegmatis. *Microbiology* **148**, 3007–3017 (2002).
